# Pulmonary B cells ameliorate Alzheimer’s disease-like neuropathology

**DOI:** 10.1101/2020.11.23.393942

**Authors:** Weixi Feng, Yanli Zhang, Tianqi Wang, Ze Wang, Yan Chen, Chengyu Sheng, Ying Zou, Yingting Pang, Junying Gao, Yongjie Zhang, Jingping Shi, Qian Li, Ming Xiao

## Abstract

Increasing evidence shows that the peripheral immune system is involved in the pathogenesis of Alzheimer’s disease (AD). Here, we report that pulmonary B cells mitigate beta-Amyloid (Aβ) pathology in 5xFAD mice. The proportion of B cells rather than T cells increases in brain, meningeal and lung tissues in 3-month-old 5xFAD mice. Deletion of B cells aggravates Aβ load and memory deficits of 5xFAD mice. Mechanimsly, pulmonary B cells can migrate to the brain parenchyma and produce interleukin-35 that inhibits neuronal β-site APP-cleaving enzyme 1 expression, subsequently reducing the production of Aβ. In turn, proliferation of pulmonary B cells is associated with activation of toll-like receptor/nuclear factor kappa-B pathway by elevated Aβ that is drained from the brain parenchyma to the lungs via meningeal lymphatics. Furthermore, promoting pulmonary B cell proliferation via overexpression of B-cell-activating factor ameliorates brain Aβ load and improves cognitive functions of 10-month-old 5xFAD mice. Together, these results highlight the lungs as both immune targets and effector organs in Aβ pathogenesis. Pulmonary B cells might be a potential target against AD.

---

Alzheimer’s disease (AD) is a neurodegenerative disease characterized by gradual accumulation of β-Amyloid (Aβ) plaques in the brain (*1*). Accumulated evidence highlights that immunity is involved in the pathogenesis of AD (*2, 3*). Previous studies mainly focus on the pathophysiological roles of microglia in AD (*4, 5*), because they serve as innate immune cells in the central nervous system (CNS). With the discovery of glymphatic system and meningeal lymphatic vessels (*6*-*8*), the contribution of the peripheral immune system in the pathogenesis of AD is gaining more attention. Here we report that meningeal lymphatics drain Aβ from the brain to the lungs, inducing B lymphocytes to produce interleukin-35 (IL-35). These pulmonary B cells then migrate to the brain and inhibit β-site APP-cleaving enzyme 1 (BACE1) expression, thus ameliorating Aβ-related pathology in 5xFAD transgenic mouse model of AD. This finding has revealed novel immunological functions of lung-brain axis in AD.

## Compensatory protections of B lymphocytes in the early stage of AD-like pathology

Adaptive immunity recently has been implicated in AD pathogenesis (*9*-*11*). For example, the number of peripheral blood B cells is positively correlated with the cognitive scores of subjects and decreases in patients with AD (*12, 13*). Loss of IgG produced by B cells impairs microglial phagocytosis, thereby exacerbating Aβ plaque deposition in Rag-5xFAD mice that lack T, B, and natural killer cells. Recently characterized lymphatic vessels in the meninges provide a crucial route for the crosstalk between the brain and peripheral immune system (*14, 15*), but their regulation in AD pathogenesis remains unknown. Based on this, we first examined changes of T and B lymphocytes in the brain parenchyma and meninges during AD-like disease progression of 5xFAD mice. Flow cytometry analysis showed that the proportion of CD19^+^ B lymphocytes increased significantly in the cortex and meninges of 5xFAD mice at age of 3 months, followed by a continuous decrease along with age (Fig. 1, A and B). However, the proportion of CD3^+^ T cells from cortex and meningeal samples was not different between 5xFAD mice and WT mice at age of 3, 5 and 8 months.

**Fig. 1.**
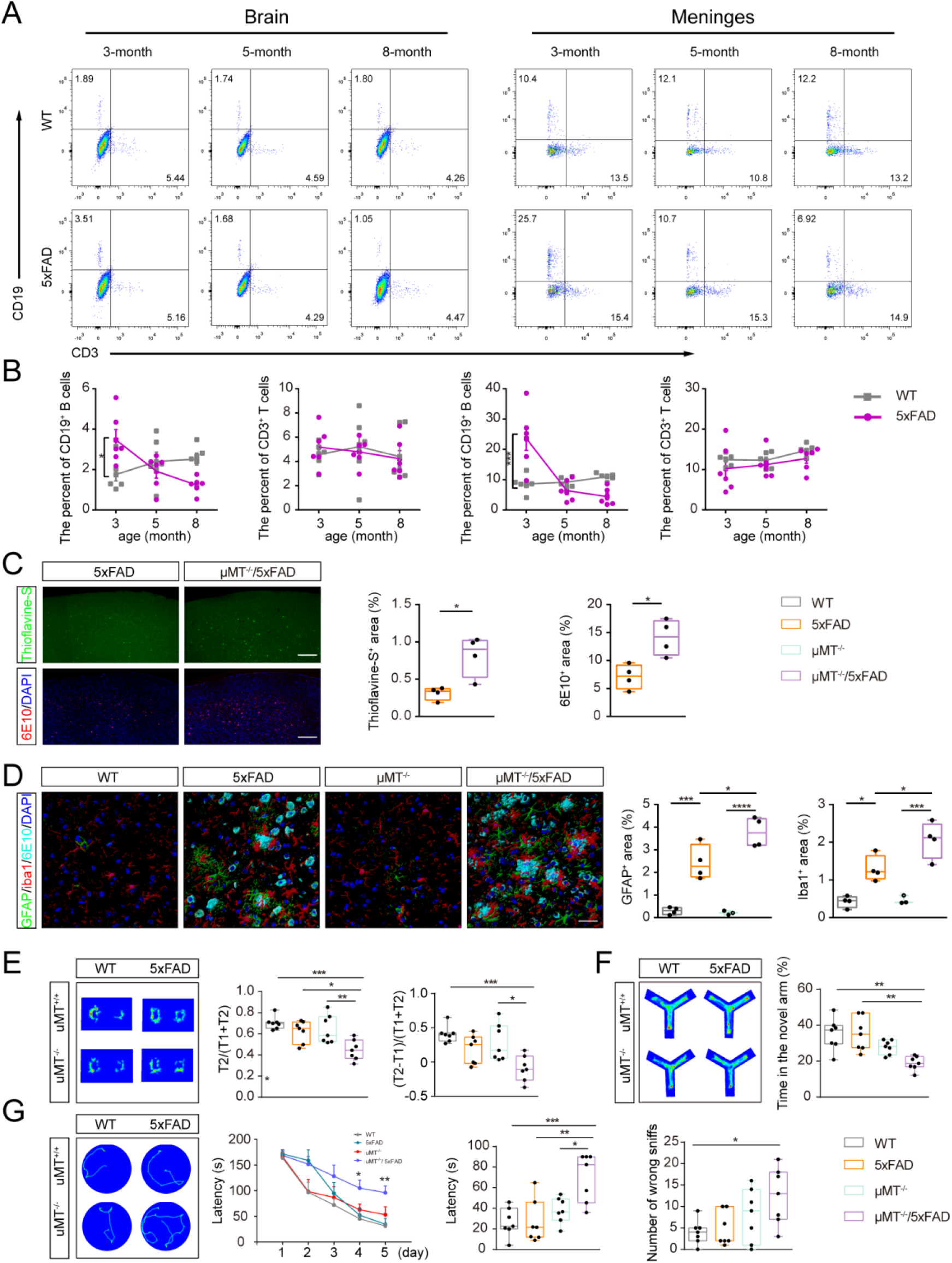
Deletion of B lymphocytes accelerated the onset of AD-like pathology in 5xFAD mice. (**A** and **B**) Representative dot plots and quantification of percentage of T cells and B cells in the brains and meninges of 3, 5 and 8 month-old WT and 5xFAD mice. n = 6 per group. (**C**) Representative images and quantification of thioflavine-S^+^ and 6E10^+^ Aβ plaques in the frontal cortex of 3-month-old 5xFAD and *μMT*^*-/-*^*/*5xFAD mice. *n* = 4 per group. (**D**) Representative images and quantification of GFAP^+^ astrocytes and Iba1^+^ microglia in the frontal cortex of 3-month-old WT, 5xFAD, *μMT*^*-/-*^ and *μMT*^*-/-*^*/*5xFAD mice. n = 3-4 per group. Scale bar = 30 μm. (**E** to **G**), Track of mice in the novel object recognition (NOR), Y maze and Barnes maze and quantification of recognition index and discrimination index in the NOR, percentage of time spent in the novel arm in Y maze and latency during the training period and test period and number of error exploration in Barnes maze of 3-month-old WT, 5xFAD, *μMT*^*-/-*^ and *μMT*^*-/-*^*/*5xFAD mice. n = 7 per group. \**P* < 0.05, \*\**P* < 0.01, \*\*\**P* < 0.001, \*\*\*\**P* < 0.0001. Data in **B, G** (latency during the training period) are analyzed by repeated-measures two-way ANOVA with Bonferroni’s post hoc test, **C** by Student’s *t*-test, the others are by one-way ANOVA with Bonferroni’s post hoc test.

In order to elucidate the role of B cells in the pathophysiology of this AD model, we crossed 5xFAD mice with B cells-deficient (*μMT*^*-/-*^) mice to generate *μMT*^*-/-*^/5xFAD mice. The pathological analysis revealed that accumulation of both thioflavine-S^+^ and 6E10^+^plaques was more obvious in the frontal cortex of three-month-old *μMT*^*-/-*^/5xFAD mice than age-matched 5xFAD littermates (Fig. 1C). *μMT*^*-/-*^/5xFAD mice showed more pronounced activation of GFAP^+^ astrocytes and Iba1^+^ microglia and significant loss of synaptic proteins synaptophysin (SYP) and postsynaptic density 95 (PSD95) in the frontal cortex (Fig. 1D and fig. S1A). Consistently, *μMT*^*-/-*^/5xFAD mice exhibited mild short-term memory impairments in both the novel objection recognition test and Y-maze test, compared with 5xFAD controls (Fig1. E and F). These mice also showed impairments in the long-term spatial learning and memory performances in the Barnes maze (Fig. 1G). These results together suggest that compensatory increases of brain B cells delay the occurrence of AD-like pathology of 5xFAD mice.

## The migration of B lymphocytes from the lungs to the brain in the early AD-like stages

Recent evidence indicates that there is a close relation between the lungs and brain for immune regulation, termed as the lung-brain axis (*16*-*18*). Moreover, the lungs might be the main target organ of central antigens drained by meningeal lymphatics (*19*). However, there is a lack of direct evidence for this hypothesis. Therefore, in order to determine the source of increased B cells in the CNS, we first detected B cells in the spleen as well as the lungs in both WT mice and 5xFAD mice. Interestingly, the proportion of CD19^+^ B lymphocytes also increased in the lungs of 3-month-old 5xFAD mice, but gradually decreased with age (Fig. 2, A to C), and there were significant positive correlations of B cells between the lungs and brain parenchyma or meninges (Fig. 2B). Nevertheless, there was no increase in the number of CD19^+^ B lymphocytes in the spleen as well as CD3^+^ T lymphocytes in the lungs or spleen at the early stages of AD-like pathology of 5xFAD mice (Fig. 2C and fig. S2, A and B).

**Fig. 2.**
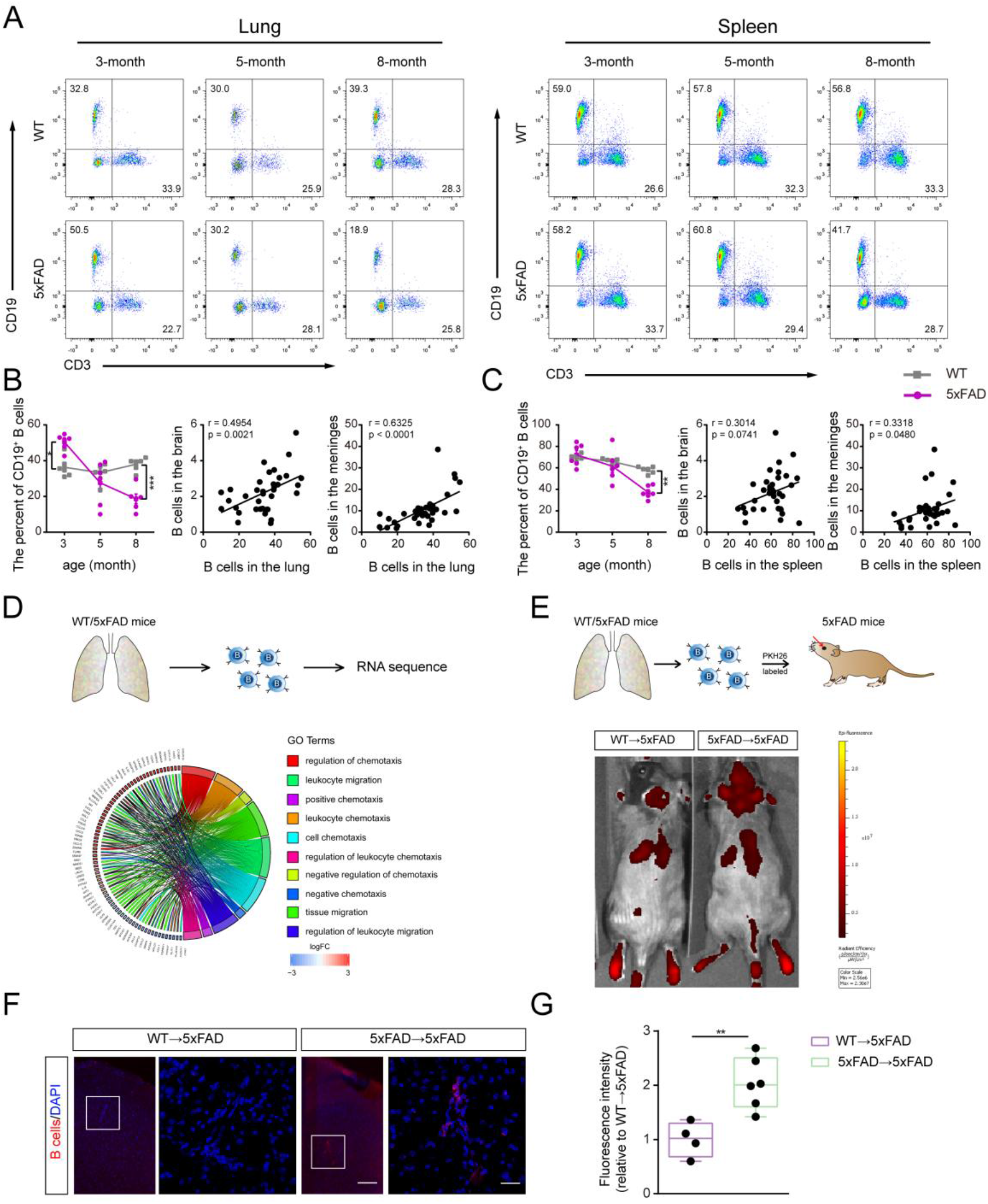
B lymphocytes migrate from the lungs to the brain in the early AD-like stages of 5xFAD mice. (**A**) Representative dot plots of T cells and B cells in the brains and meninges of 3, 5 and 8 month-old WT and 5xFAD mice. (**B** and **C**) Percentage of B lymphocytes in the lungs (B) or the spleen (C) of 3-, 5- and 8-month-old WT mice and 5xFAD mice, and correlation analysis of the percentage of B lymphocytes between the lungs and the brains or meninges respectively, n = 6 per group. (**D**) Gene Ontology (GO) analysis of B lymphocytes mRNA-seq in the lungs of 3-month-old WT and 5xFAD mice. (**E**) PKH26 labeled pulmonary B lymphocytes tracking to brain. (**F**) Labeled B cells observed in the frontal cortex by immunofluorescence. (**G**) Fluorescence intensity of labeled B cells in the frontal cortex. n = 5-6 per group. Scale bar = 30 μm. \**P* < 0.05, \*\**P* < 0.01, \*\*\**P* < 0.001, \*\*\*\**P* < 0.0001. Data represents as mean ± s.e.m. Percentage of B lymphocytes in **B, C** are analyzed by repeated-measures two-way ANOVA with Bonferroni’s post hoc test, and correlation analysis in **B, C** by Pearson Correlation Coefficient and data in **G** by Student’s *t*-test.

In order to elucidate how these lung-derived B cells change along with AD-like disease progression, we first conducted RNA-sequencing on sorted B cells from the lungs of 3-month-old 5xFAD mice and WT mice. Gene Ontology (GO) analysis showed that differentially expressed genes between the two groups were significantly enriched in cell migration (Fig. 2D and fig. S2C). We then conducted B cell transplantation experiments to exanmin whether lung-derived B cells can migrate to the brain. B cells from the lung of 3-month-old WT or 5xFAD mice were infused into 5xFAD mice with the same age via retro-orbital injection. The *in vivo* fluorescent imaging demonstrated that more B cells from 5xFAD lungs were detected in the brain at 24 h after transplantation compared with those from WT mice (Fig. 2, E and G). Immunofluorescent results further confirmed transplanted B cells within the cortex of 5xFAD mice (Fig. 2F). Together, these results indicate that B cells from 5xFAD mice lungs migrate efficiently into the brain, which contributes to the increase of B cells in the early stages of amyloidogenesis.

## IL-35 mediates pulmonary B lymphocytes against Aβ pathology in 5xFAD mice

In order to investigate the mechanism underlying the alleviating effect of lung-derived B cells on the onset of AD-like pathology, we further compared the differentially expressed genes of pulmonary B cells between 3-month-old WT mice and 5xFAD mice. GO analysis revealed that differentially expressed genes were also significantly enriched in the categories of immune activation, cytokine production and secretion (Fig. 3, A to C). Notably, we found that Epstein-Barr virus-induced gene 3 (Ebi3), one of the subunits of IL-35, was upregulated in sorted B cells from the lungs of 3-month-old 5xFAD mice (Fig. 3C). IL-35, as a new cytokine, has been shown to regulate immune response and participate in a variety of pathological conditions including the central nervous system (CNS) disorders (*20*-*23*). For example, B-cell with IL-35 deficiency aggravates demyelination in an experimental autoimmune encephalomyelitis (EAE) mouse model (*22*). However, it is not known whether IL-35 participates in AD pathogenesis. Therefore, we focused on the role of pulmonary B cells-derived IL-35 in Aβ pathology in 5xFAD mice.

**Fig. 3.**
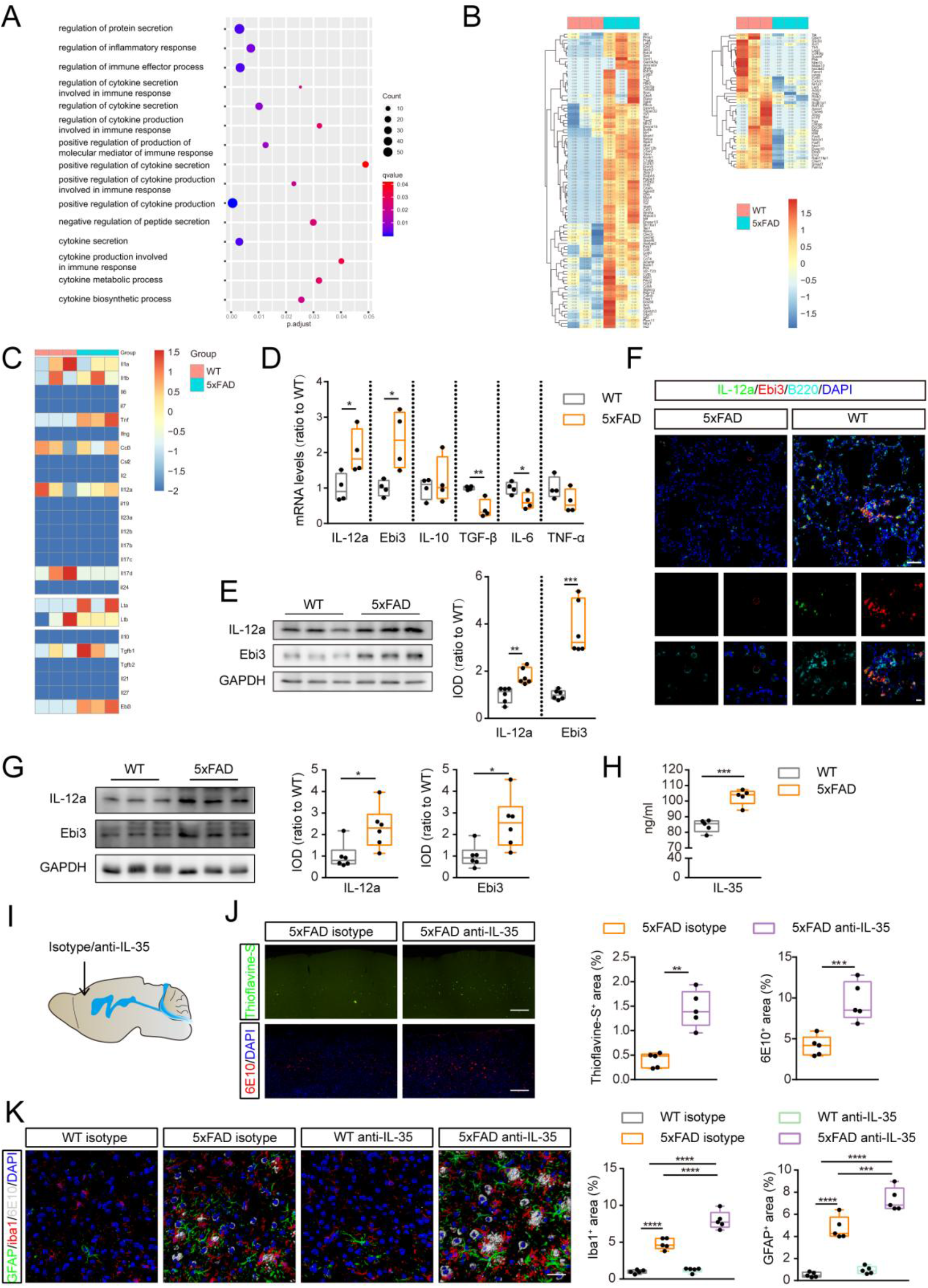
B cells increase IL-35 secretion in the early AD-like stages of 5xFAD mice. (**A**) GO analysis of B cells mRNA-seq in the lungs of 3-month-old WT and 5xFAD mice. n = 3 per group. (**B** and **C**) Heat map showing relative expression level of genes involved in mediating protein secretion and cytokines. Color scale bar values represent standardized log-transformed values across samples. n = 3 per group. (**D**) QRT-PCR for *IL-12a, Ebi3, IL-10, TGF-β, IL-6* and *TNF-α* of B cells in the lungs. n = 4 per group. (**E**) Western blot for IL-12a and Ebi3 in the lungs of 3-month-old mice. n = 6 per group. (**F**) B cells in the lungs with high expression of IL-12a and Ebi3 in 5xFAD mice. Scale bar = 50 and 10 μm. (**G**) Western blot for IL-12a and Ebi3 in the cortex of 3-month-old mice. n = 6 per group. (**H**) The level of IL-35 in the frontal cortex detected by ELISA. n = 5 per group. (**I** and **J**) Representative images and quantification of thioflavine-S^+^ and 6E10^+^ plaques in the frontal cortex of 3-month-old 5xFAD with IL-35 neutralization or not. n = 5 per group. Scale bar = 30 μm. (**K**) Representative images and quantification of GFAP^+^ astrocytes and iba-1^+^ microglia in the frontal cortex of 3-month-old 5xFAD with IL-35 neutralization or not. n = 5 per group. Scale bar = 30 μm. Data represents as mean ± s.e.m. * *P* < 0.05, \*\**P* < 0.01, \*\*\**P* < 0.001, \*\*\*\**P* < 0.0001. Data in **B-J** are analyzed by Student’s *t*-test, and in **K** by ANOVA with Bonferroni’s post hoc test.

Both mRNA and protein levels of IL-12a and Ebi3, two subunits of IL-35, increased in the lungs of 3-month-old 5xFAD mice (Fig. 3, D and E). Immunostaining results demonstrated that IL-12a^+^Ebi3^+^ B cells were accumulated in lung tissues of 3-month-old 5xFAD mice (Fig. 3F). Consistent with the increase in the proportion of B cells in the brain parenchyma, western blot and ELISA analysis confirmed that IL-12a and Ebi3 levels in the frontal cortex were significantly higher in 3-month-old 5xFAD mice than WT mice (Fig. 3, G and H). B cells deficiency blocked the up-regulation of IL-35 expression in the frontal cortex of 3-month-old 5xFAD mice (fig. S3A), suggesting an important role of B cells in IL-35 secretion. In order to determine whether the increased IL-35 alleviates Aβ-related pathology, we neutralized IL-35 in the frontal cortex of 3-month-old 5xFAD mice by stereotactic injection of IL-35 antibody. Five days after injection, histopathological and biochemical analyses showed that IL-35 neutralization significantly increased intraneuronal Aβ aggregation and extracellular plaque disposition, evoked activation of astrocytes and microglia, and exacerbated the loss of SYP in the frontal cortex of 5xFAD mice (Fig. 3, I to K and fig. S3B).

## IL-35 inhibits BACE1 expression in neurons via SOCS1/STAT1 pathway

Aβ accumulation within the brain is attributed to an imbalance between its production and clearance (*24*). In order to investigate the potential mechanisms of IL-35 reducing brain Aβ load in 5xFAD mice, we examined the expression of several enzymes or proteins involved in brain Aβ metabolism. Western blot results showed that neutralization of IL-35 in the frontal cortex of 3-month-old 5xFAD mice neither affected the expression of Aβ precursor protein (APP) secretases A disintegrin and metalloproteinase domain-containing protein 10 (ADAM10) and presenilin-1 (PS1) (fig. S3C), nor changed Aβ clearance-related markers including neprilysin (NEP), insulin-degrading enzyme (IDE) and low-density lipoprotein receptor-related protein 1 (LRP1) (fig. S3D). However, the expression levels of BACE1, a key amyloidogenic enzyme, significantly increased in the 5xFAD frontal cortex after IL-35 neutralization (Fig. 4A). Similar changes were also observed in the cortex of 3-month-old *μMT*^*-/-*^/5xFAD mice when compared with 5xFAD controls (fig. S4A). We further investigated whether elevated BACE1 affects APP hydrolysis process. Western blot results showed that IL-35 neutralization or B cell deficiency significantly increased amyloidogenic products soluble amyloid precursor protein β (sAPPβ) and C-terminal fragment β (CTFβ) in the frontal cortex of 5xFAD mice, without affecting the levels of APP as well as non-amyloidogenic products soluble amyloid precursor protein α (sAPPα) and C-terminal fragment α (CTFα) (Fig. 4A and fig. S4B). These data suggested that IL-35 inhibits BACE1 expression, subsequently reducing amyloidogenic products.

**Fig. 4.**
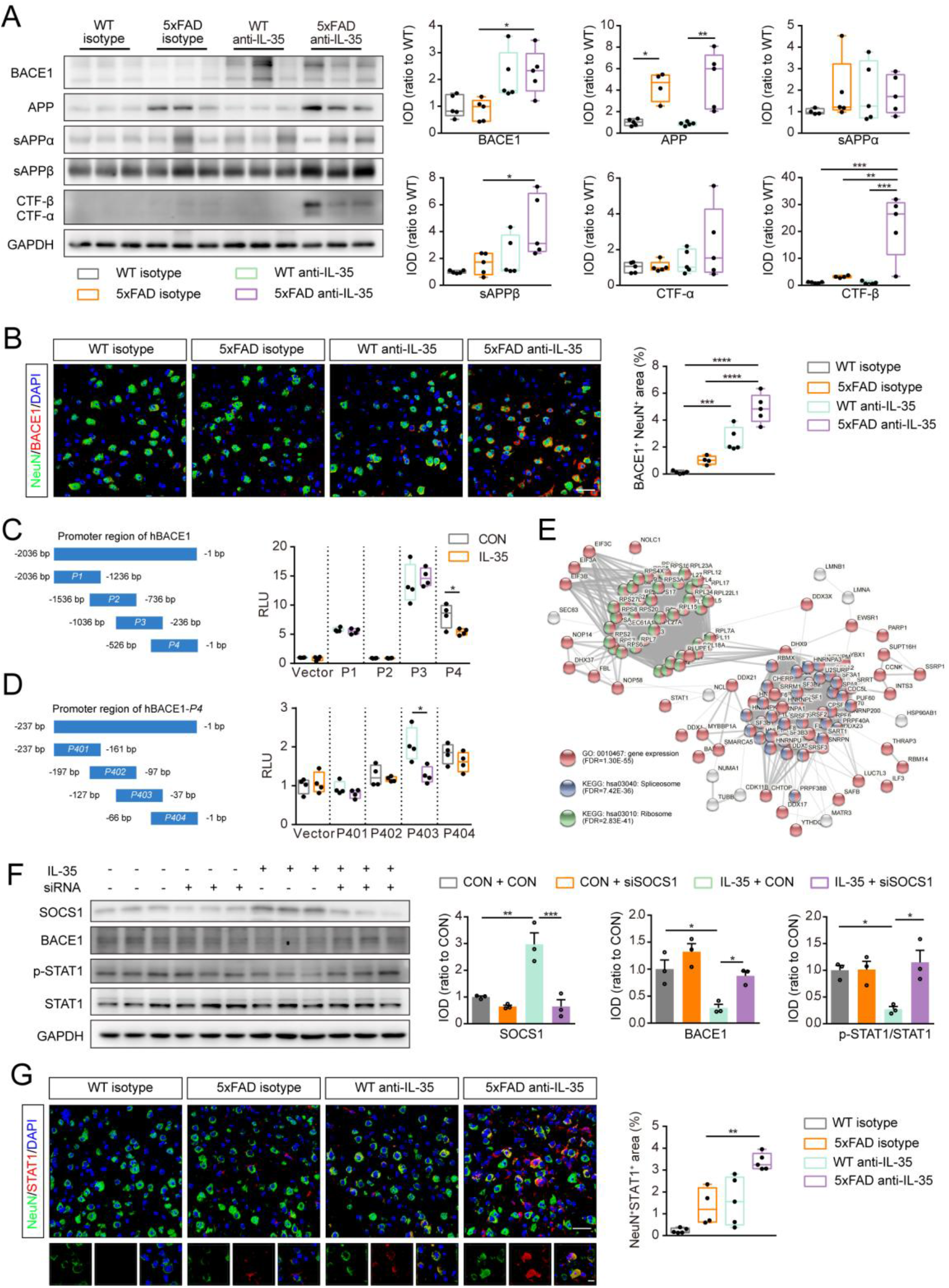
IL-35 inhibits BACE1 expression in neurons via SOCS1/STAT1 pathway. (**A**) Representative Western blot bands and quantification of APP secretases BACE1 and APP and its main cleaved fragments sAPPα, sAPPβ, CTF-α and CTF-β in the PFC. n = 5 per group. (**B**) Representative images of NeuN with BACE1 in the PFC and quantification of BACE1^+^ areas in neurons after IL-35 neutralized. n = 5 per group. Scale bar = 30 μm. (**C and D**) Luciferase assay for BACE1 transcription activity after IL-35 treatment. n = 4 per group. (**E**) String interaction prediction was performed (https://string-db.org/) indicating interactions between spliceosome and ribosome proteins. (**F**) Representative bands and quantification of Western blot for SOCS1, BACE1, p-STAT1 and STAT1 after knocking-down of SOCS1 expression or IL-35 treatment in SH-SY5Y cells. n = 3 per group. (**G**) Representative images and quantification of NeuN and STAT1 after IL-35 neutralization. n = 5 per group. Scale bar = 30 and 10 μm. \**P* < 0.05, \*\**P* < 0.01, \*\*\**P* < 0.001, \*\*\*\**P* < 0.0001. Data in **C, D** analyzed by Student’s t-test, **A, B, F, G** by ANOVA with Bonferroni’s post hoc test.

Previous studies reported that IL-12RB2 and GP130 form homodimers or heterodimers that act as IL-35 cell surface receptors to conduct intracellular downstream signals (*25*). IL-35 receptors were localized to prefrontal cortical neurons, as revealed by double-immunofluorescence for IL-12RB2 or GP130 with NeuN, respectively (fig. S5A). Furthermore, we demonstrated the existence of IL-35 and its receptors in the postmortem cortex (fig. S6). Notably, cortical neurons increased BACE1 expression and 6E10^+^ plaque subsequently after local neutralization of IL-35 (Fig. 4B and fig. S5B). *In vitro* experiments also showed that BACE1 expression was reduced in mouse primary cortical neurons, mouse neuron cell line N2a and human neuron cell line SH-SY5Y treated with IL-35 recombinant protein for 48 h, without affecting ADAM10 and PS1 levels (fig. S5, C to H). Furthermore, BACE1 enzyme activity was significantly reduced after IL-35 stimulation in primary cortical neurons (fig. S5I). These results indicated that IL-35 inhibits neuronal BACE1 expression via its receptors.

Next, we investigated how IL-35 regulates the expression of BACE1. The -2036 to -1 bp region upstream from the transcription start site (TSS) was divided into four segments (P1-P4) to make partially overlapping reporter constructs, each 526 to 800 bp in length. Dual-luciferase assay revealed that IL-35 significantly down-regulated hBACE1 promoter segments P4 activity (Fig. 4C). We further divided P4 into four segments, namely P401 to P404 encompassed only the 30 bp region. The results revealed that IL-35 regulated BACE1 transcriptional level through P403 (−127bp to -37bp) region (Fig. 4D). Then, we identified the proteins binding P403 by DNA affinity chromatography pull-down and mass spectrometry assay to elucidate the transcription factors involved in the regulation of BACE1. The proteins showing significant interaction with the proximal promoter are listed in Supplementary Table 1. In addition, these analyses also identified clusters of spliceosome and ribosome proteins (RBPs) that strongly interacted with the aforementioned promoter segment (Fig. 4E). Interestingly, transcription factor signal transducer and activator of transcription 1 (STAT1) was also observed in the binding proteins and showed interaction with the spliceosome and ribosome clusters (Fig. 4E). Previous studies have revealed that STAT1 is a key transcription factor for BACE1 in neurons (*26*). We further confirmed the binding of STAT1 to hBACE1 P403 fragment by Western blot (fig. S5J). We also examined another BACE1 transcription factor, c-jun (*26*), which was not detected in the binding proteins (fig. S5J). Meanwhile, suppression of STAT1 activation is mainly mediated by suppressor of cytokine signaling (SOCS), which plays an important role in the regulation of intracellular inflammatory signals (*27*). Based on this, we tested the effect of IL-35 on SOCS/STAT1 signaling pathway in primary mouse neurons, N2a cells and SH-SY5Y cells. IL-35 stimulation significantly increased the expression of SOCS1 rather than SOCS3 in the above three types of cultured cells (fig. S5, C to H). Consistently, knockdown of SOCS1 reversed the inhibitory effect of IL-35 on BACE1 expression and STAT1 phosphorylation in SH-SY5Y cells (Fig. 4F). SOCS1 protein levels and STAT1 phosphorylation also significantly increased after IL-35 neutralization in the frontal cortex of 3-month-old 5xFAD mice (fig. S5K). Immunofluorescence staining further showed that IL-35 neutralization resulted in an increase of STAT1 expression in neuronal nucleus of 5xFAD mice (Fig. 4G). The above results together indicated that IL-35 inhibited the transcription of BACE1 via SOCS1/STAT1 signaling pathway.

## Meningeal lymphatics drainage Aβ that promotes B lymphocytes to produce IL-35 via TLR/NF-κB pathway

Typical lymphatic vessels have been identified in the human and mouse dura recently (*6, 7*), and serve as an important drainage route for CNS macromolecules and antigens (*3, 14*). There is also the glymphatic pathway that transports interstitial metabolites from the brain parenchyma to the subarachnoid space (*8, 28*). Blocking meningeal lymphatic vessels or ligating the deep cervical lymph nodes (dcLNs) impairs glymphatic transport, resulting in an increase of Aβ load (*3, 29*). These findings suggest that the glymphatic and meningeal lymphatic systems constitute a functional macromolecular drainage pathway. However, the peripheral target organs and biological effects of these CNS antigens have yet to be fully determined. Therefore, we injected Evans blue into the cistern magna and found the dcLNs were stained by Evans blue only from 8 min after injection. At the same time, a gradual darkening of the dye was observed in the lungs (Fig. 5A). Nevertheless, there was less Evans blue content in the spleen at 8 min after injection (Fig. 5B). We further determined whether ligation of the dcLNs would affect brain Aβ to reach the lungs. As expected, dcLNs ligation reduced the content of cortical exogenous injected AF555-labeled Aβ in the lungs (Fig. 5, C and D). Furthermore, endogenous oligo-Aβ levels were significantly reduced in the lung tissues of 3 month-old 5xFAD mice 1 month after dcLN ligation, without changes in the APP content (Fig. 5, E to G). These results have revealed that macromolecules including Aβ drained by meningeal lymphatic vessels could be transported from the brain parenchyma to the dcLNs, enter the superior vena cava, and finally reach the lungs via the pulmonary circulation. We also found that ligation of the dcLNs significantly decreased the content of other specific central antigens including microtubule associated protein 2 (MAP2) in the lung tissues (fig. S7A).

**Fig. 5.**
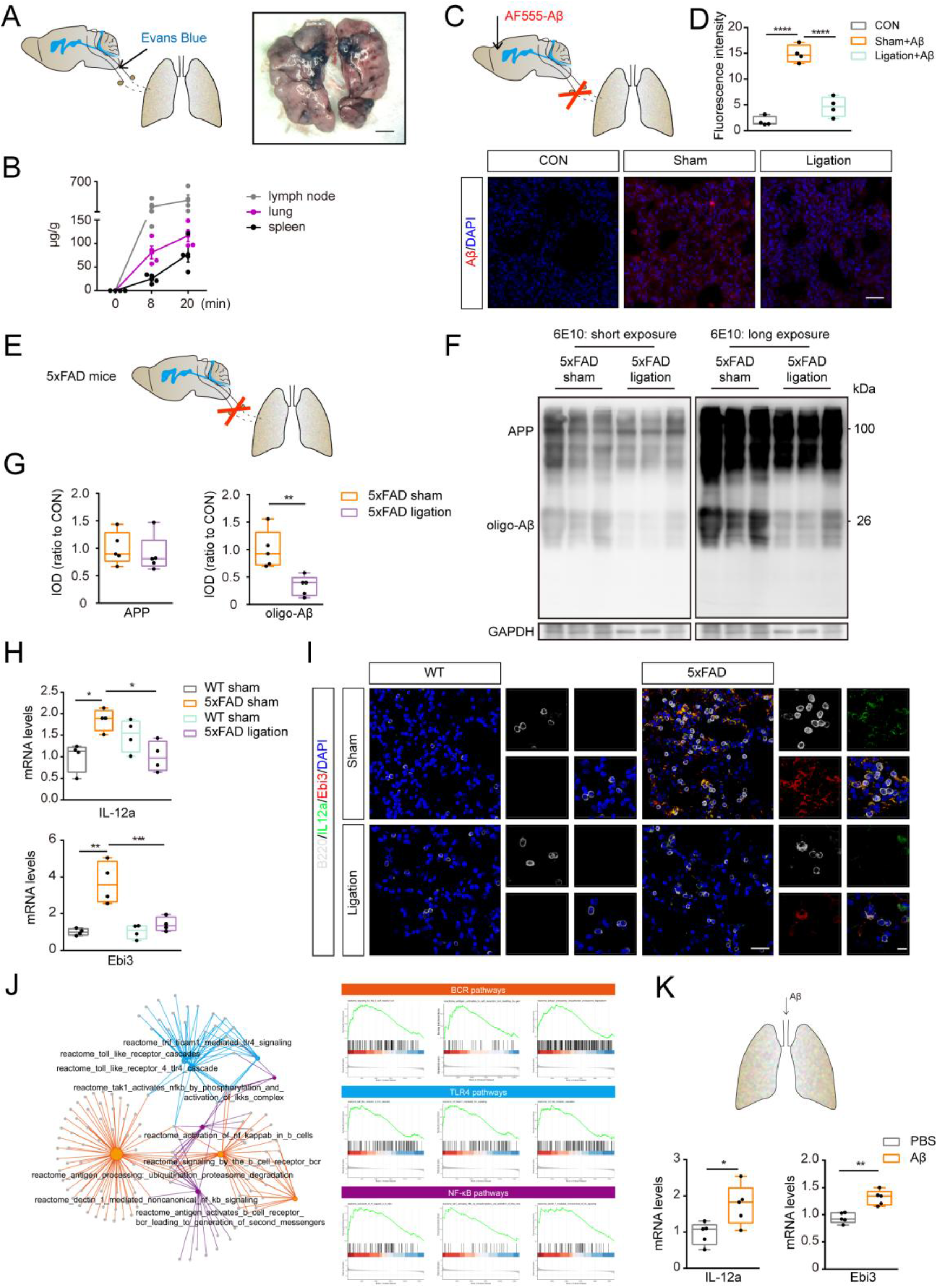
Aβ drainage from meningeal lymphatic promotes IL-35 secretion of B lymphocytes in the lungs. (**A** and **B**) Representative image of Evans blue in the lungs and quantification of Evans blue in the dcLNs, lungs and spleen after injection. n = 4 per group. Scale bar = 30 μm. (**C** and **D**) Representative images and quantification of AF555-labeled Aβ in the lungs after dcLN ligation. n = 4 per groups. Scale bar = 30 μm. (**E** to **G**) Representative bands and quantification for APP and Aβ in the lungs of 3-month-old mice after dcLN ligation. n = 5 per group. (**H**) QRT-PCR for *il12a* and *ebi3* after dcLN ligation. n = 4 per group. (**I**) Representative images for the expression of IL-12a and Ebi3 in B cells in the lungs of 3-month-old mice after dcLN ligation. Scale bar = 30 and 10 μm. (**J**) GSEA analysis of B lymphocyte gene expression in the lungs of 3-month-old mice showed that BCR pathways, TLR4 pathways and NF-κB pathways were activated in the B cells of 5xFAD mice. (**K**) mRNA level of IL-12a and Ebi3 in the pulmonary B cells after Aβ treatment. n = 5 per group. \**P* < 0.05, \*\**P* < 0.01, \*\*\**P* < 0.001, \*\*\*\**P* < 0.0001. Data in **G** and **K** analyzed by Student’s t-test, **B** by repeated-measures ANOVA with Bonferroni’s post hoc test, **D, H** by ANOVA with Bonferroni’s post hoc test.

Moreover, blocking meningeal lymphatic drainage by ligation of the dcLNs reduced the secretion of IL-35 by pulmonary B cells in 3-month-old 5xFAD mice (Fig. 5, H and I). Brain Aβ can induce the phenotype changes of microglia via toll-like receptors (TLR) signaling pathway (*30*). In order to determine whether increased IL-35 production by pulmonary B cells is also related to Aβ, we performed GSEA analysis on pulmonary B cell transcriptome of 3-month-old WT and 5xFAD mice. The results demonstrated that the genes associated with TLR, B-cell receptor (BCR), and nuclear factor-kappa B (NF-κB) pathways were up-regulated significantly in 5xFAD mice, compared to age-matched WT littermates (Fig. 5J), which were consistent with previous reports that activation of TLR and BCR participates in differentiation of B cell and IL-35 secretion (*31*). Interestingly, Aβ has been reported to activate TLR and NF-κB pathways (*32*), suggesting that Aβ may contribute to IL-35 secretion from the lung-derived B cells in the early AD-like stage of 5xFAD mice. In order to further verify this hypothesis, hAβ_1-42_ peptides were injected intratracheally to the lungs of three-month-old WT mice. The results showed that the mRNA levels of IL-12a and Ebi3 increased significantly in the pulmonary B cells 5 days after injection (Fig. 5K). Consistently, TLR and NF-κB pathways associated genes markedly up-regulated as well (fig. S7B). Together, these results suggested that the lungs not only received CNS lymphatic draining Aβ, but also served as an immune effector organ of Aβ.

## B-cell activation factor (BAFF) overexpression induces pulmonary B lymphocyte proliferation alleviates Aβ pathology in 5xFAD mice

Previous studies have demonstrated that BAFF promotes the survival of B cells and induces IL-35 secretion (*33, 34*). The above results showed that the secretion of IL-35 in B cells induced by Aβ could inhibit BACE1 expression and ameliorate Aβ pathology subsequently. To explore the therapeutic potential of regulating IL-35 secretion from lung-derived B cells against Aβ pathology, we overexpressed BAFF in the lungs of 7-month-old 5xFAD mice to induce the increase of B cells in the lungs and the brain (Fig. 6, A and B). The mRNA levels of IL-12a and Ebi3 were up-regulated in the lungs of 10-month-old 5xFAD mice after overexpression of BAFF for 3 months (Fig. 6C). The pathological analysis revealed that overexpression of BAFF in 5xFAD mice inhibited the expression of BACE1 and reduced 6E10^+^ plaque load in the frontal cortex (Fig. 6, C and D). Consistently, injection of AAV-BAFF improved cognitive functions of 5xFAD mice, revealed by increased residence time in the novel arm in the Y-maze and decreased latency in the Barnes maze (Fig. 6, E to G). The above results have demonstrated that the pulmonary BAFF overexpression can mitigate Aβ load and improve cognitive functions of 5xFAD mice even at the mid-late AD-like stages.

**Fig. 6.**
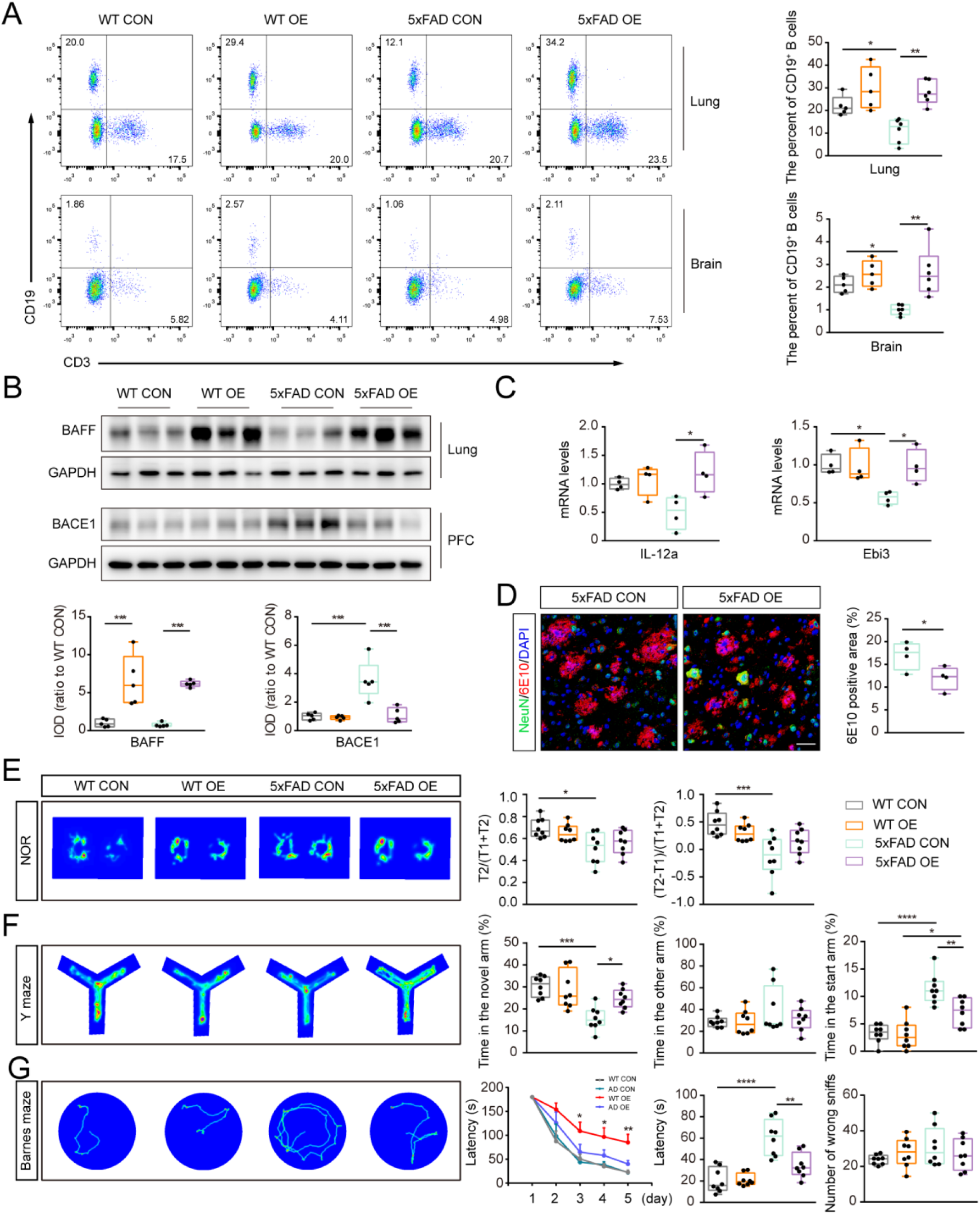
Over-expression of BAFF in the lungs ameliorates memory impairment in the lungs of 10-month-old 5xFAD mice. (**A**) Representative dot plot and quantification of B lymphocytes populations in the lungs and brain of 10-month-old mice after AAV-mBAFF injection. n = 5 per group. (**B**) Representative bands and quantification of BAFF in the lungs and BCAE1 in the frontal cortex of 8-month-old WT and 5xFAD mice after AAV-mBAFF injection. n = 5 per groups. (**C**) Quantification of mRNA levels of IL-12a and Ebi3 in pulmonary B cells after AAV-mBAFF injection. n = 4 per group. (**D**) Representation images and quantification of 6E10 and NeuN in the frontal cortex of 10-month-old 5xFAD mice after AAV-mBAFF injection. n = 4 per groups. Scale bar = 30 μm. (**E**) Track plot of novel object recognition in 10-month-old mice and analysis of recognition index and discrimination index. (**F**) Track plot for 10-month-old mice in the Y maze test and percentage of time taken in the novel arm. (**G**) Track plot of the Barnes maze test in the 10-month-old mice and statistics of incubation period of training, incubation period of test and number of error exploration. n = 8 per group.\**P* < 0.05, \*\**P* < 0.01, \*\*\**P* < 0.001, \*\*\*\**P* < 0.0001. Data are mean ± s.e.m. **A** to **C** by ANOVA with Bonferroni’s post hoc test. **D** analyzed by Student’s test. **E** to **G (**latency during the training period) are analyzed by repeated-measures two-way ANOVA with Bonferroni’s post hoc test, the others are by one-way ANOVA with Bonferroni’s post hoc test.

## Discussion

Taken together, the present findings highlight potential for pulmonary B cells against Aβ pathology. The number of B cells increases temporarily in the brain, meninges and lungs in the early stage of AD-like neuropathology in 5xFAD mice. Deletion of B cells impairs the short-term and long-term memory of 3-month-old 5xFAD mice, accompanied with more severe Aβ pathology. We further found lung-derived B cells can migrate to the brain parenchyma and produce IL-35 that inhibits neuronal BACE1 expression through the SOCS1/STAT1 pathway, subsequently reducing the production of Aβ. In turn, proliferation of pulmonary B cells is associated with activation of TLR/NF-κB pathway by elevated Aβ, which is drained from the brain parenchyma to the lungs via meningeal lymphatics. Blocking meningeal lymphatic drainage markedly reduces Aβ levels and B cell number in the lungs. Furthermore, promoting pulmonary B cell proliferation and IL-35 secretion via overexpression of BAFF ameliorates AD-like pathology in 5xFAD mice. These results collectively suggest that the lungs serve as sites for B cell activation in response to brain Aβ accumulation and migration into the brain to inhibit Aβ production via IL-35/SOCS1/BACE1 pathway. However, the mechanism of reduced production of lymphocyte B and IL-35 in the late stages of AD-like disease progression still needs further elucidation.

Cross-talks between the lungs and brain have been documented recently (*16*-*18*). For example, activated T cells in the experimental autoimmune encephalomyelitis model are observed to be enriched in the lungs before invading the brain parenchyma where they up-regulate the expression of adhesion molecules and chemokines (*16*). Nevertheless, disruption of meningeal lymphatic drainage diminishes pathology and reduces the inflammatory response of brain-reactive T cells during an animal model of multiple sclerosis (*14*). However, the anatomical basis for the brain-lung axis remains elusive. The present results have revealed this “black box”. Aβ and a variety of central antigens can reach the lungs through meningeal lymphatic vessels. Recently, the role of B cells in neuroinflammation and meningeal immunity has gradually gained a new understanding (*15, 35*). This study further indicates that decreased pulmonary B cells infiltrating brain also partially contributes to increased Aβ load following blocking meningeal lymphatic drainage.

Inhibition of BACE1 expression and activity to reduce Aβ load in the brain has been considered as a key target for AD therapy. However, the safety and side effects of small molecule inhibitors of BACE1 are also of concern (*36*). In the present study, we have identified a natural signaling pathway that specifically inhibits BACE1 expression by inducing the expression of IL-35 in pulmonary B cells. Notably, IL-35, as an anti-inflammatory factor, can also directly inhibit neuroinflammation in AD brain (*22, 25*). Therefore, regulation of pulmonary B cells to produce IL-35 may provide a new avenue against AD, but its clinical efficacy and safety need to be further defined.

## Author contributions

W.F., Y.Z., T.W. Z.W., Y.C. conducted the experiments. W.F., Y.Z., Y.Z., Y.P., J.G. carried out the data analysis. M.X., W.F., Q.L., C.S., J.S., Y.Z. designed the experiments. M.X., W.F., Q.L. wrote the manuscript. All authors read and approved the final manuscript.

## Data and materials availability

Supplementary materials contain additional data. All data needed to evaluate the conclusions in the paper are present in the paper or the supplementary materials. RNA-sequencing data are available through GEO no (under application).

## Schema

An involvement of “lung-brain” axis in the Aβ related pathology. Brain derived Aβ are transported to the lungs in whch they activates IL-35-produced B cells. These B cells migrate to the brain and down-regulate neuronal BACE1 expression via IL-35/SOCS1/STAT1 pathway, thus inhibiting Aβ production subsequently.

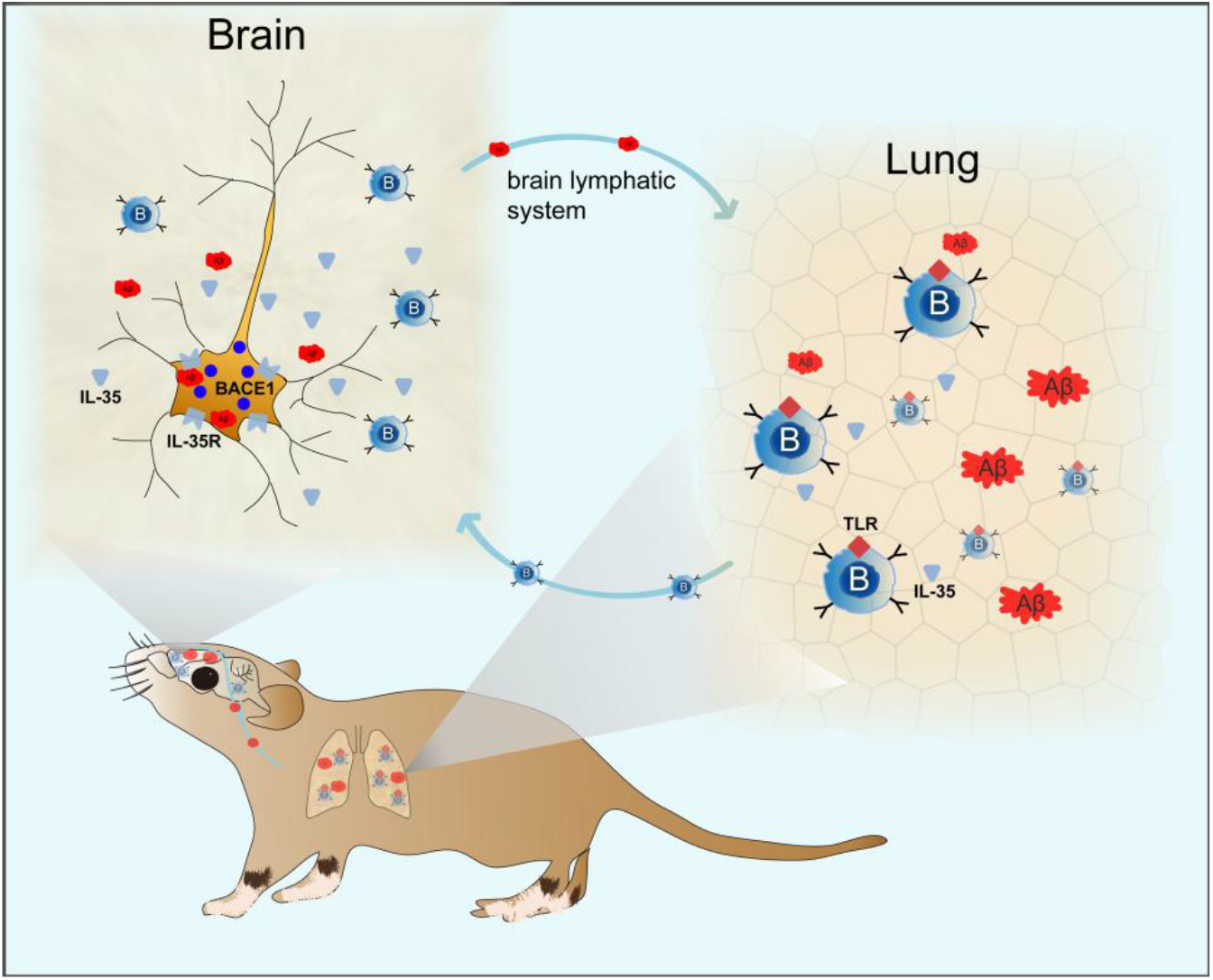

