## Supplementary material for "Pulmonary B cells ameliorate Alzheimer’s disease-like neuropathology": Methods and supplemental figures and tables

**Mouse strains and housing**

Male wild-type mice (C57BL/6J background) and B6.Cg-Tg (*APPSwFlLon, PSEN1*M146L*L286V*) 6799Vas/Mmjax (5xFAD, JAX 008730) were purchased from the Jackson Laboratory. *μMT*$^{-/-}$ mice which are deficient in B cells were purchased from Shanghai Model Organisms Center, Inc. Both B6.Cg-Tg and *μMT*$^{-/-}$ mice were bred on a C57BL/6J background. 5xFAD mice and *μMT*$^{-/-}$ mice were crossed to generate *μMT*$^{-/-}$/5xFAD mice. The mice were used at different ages that are indicated throughout the manuscript. Mice of all strains were specific pathogen free environment with controlled temperature and humidity, on 12 h light: dark cycles (lights on at 7:00), and fed with regular rodent's chow and sterilized tap water ad libitum. All experiments were approved by the Institutional Animal Care and Use Committee of the Nanjing Medical University.

**Human samples**

Frontal cortex tissues were obtained from 4 cases within 4-6 h of death, via informed donation for the Medical Education and Research of Nanjing Medical University, with corresponding written consents prepared by the donors and their families. Cases that were died from brain associated diseases were excluded from this study. The utilization of human tissues was approved by the Ethics Committee of Nanjing Medical University. All obtained samples were fixed in a 4% formalin solution and kept in paraffin blocks until further sectioning.

**Animal surgery**

**DcLN ligation:** The procedure of surgical ligation of the lymphatics afferent to the dcLNs was according to published literature (*3, 30*). In brief, mice were anaesthetized by i.p. injection with ketamine and xylazine in saline, and fixed on a stereotaxic apparatus in a supine position. The skin of the neck was shaved and cleaned with iodine and 70% ethanol and ophthalmic solution placed on the eyes to prevent drying. A midline incision was made 5 mm superior to the clavicle. Fat and soft tissues were separated with a blunt forceps under the microscope. Then, the sternocleidomastoid muscles were retracted and both sides of the dcLNs were exposed. Their afferent vessels were carefully ligated using 8-0 nylon suture. The mouse skin was sutured and disinfected with iodophor. Control mice were subjected to a sham surgery consisting of the skin incision and retraction of the sternocleidomastoid muscle only. The skin was then sutured, after which the mice were subcutaneously injected with ketoprofen (2 mg $kg^{-1}$) and allowed to recover on a heat pad until fully awake.

**Intratracheal injection:** Mice were anaesthetized by i.p. injection of ketamine and xylazine in saline and fixed on a stereotaxic apparatus in a supine position. The skin of the neck was shaved and cleaned with iodine and 70% ethanol and ophthalmic solution placed on the eyes to prevent drying. A midline incision was made 5 mm superior to the clavicle. A rubber duct connected with a 50 μL syringe was inserted into trachea through a small incision bellow the cricoid cartilage. The injection was finished within 5 min followed by an additional administration of 200 mL air to push the liquid into the bronchi. For AAV delivery, WT or 5xFAD mice were injected with $10^{11}$ of AAV5 encoding mBAFF under CMV promoter or control virus in 20 μL volume at the age of 7 months old and sacrificed when they were 10-month-old. For Aβ delivery, 3-month-old WT mice were injected with 20 μg of $A\beta_{1\text{-}42}$ in 20 μL volume and sacrificed 5 days later.

**Brain parenchymal injection:** Mice were anaesthetized by i.p. injection of ketamine and xylazine in saline and the head was secured in a stereotaxic frame. An incision was made in the skin to expose the skull. For IL-35 neutralization experiments, 1.5 μL of anti-Ebi3 antibody (ROCKLAND, 210-301-B66) or isotype control antibody was injected to mice frontal cortex (anteroposterior: − 2.5 mm; mediolateral: ±1.0 mm; dorsoventral: − 0.5 mm relative to bregma) using a Hamilton syringe (coupled to a 33-gauge needle) at a rate of 0.3 μL/min. For Aβ tracking experiments, 2 μL of AF555-Aβ or PBS control was injected to mouse frontal cortex (anteroposterior: − 2.5 mm; mediolateral: ±1.0 mm; dorsoventral: − 0.5 mm relative to bregma) using a Hamilton syringe (coupled to a 33-gauge needle) at a rate of 0.3 μL/min. After injecting, the syringe was left in place for additional 5 min to prevent backflow. The scalp skin was then sutured, after which the mice were subcutaneously injected with ketoprofen (2 mg $kg^{-1}$) and allowed to recover on a heat pad until fully awake.

**Behavioral test**

**Novel objection recognition test:** The novel objection recognition test was performed following a published protocol. The experimental apparatus used in this study was rectangular box made of opaque white plastic (50 cm × 35 cm). The mice were first habituated to the apparatus for 5 min. Two same plastic objects were then positioned in two side of the box and 5 cm away from each adjacent arena wall. Mice were then placed in the arena and allowed to explore the arena and objects for 5 min. After 2 h, the mice were placed in the same box with the two objects for another 5 min, but one of them had been replaced with a new objects in different shapes. Exploration of an object was assumed when the mouse approached an object and touched it with its vibrissae, snout or forepaws and was measured using a video tracking software (TopScan, CleverSys, Inc.). T1 represented the exploration time for new object and T2 for the other object. The recognition index was calculated as T2/(T1+T1) and the discrimination index was calculated as (T2-T1)/(T1+T2).

**Barnes maze test** (BM): The BM test was performed as described previously (*37*), with minor modifications. Mice were transported to the behavior room to habituate at least 2 hours before starting the test. The BM test consisted of five days of acquisition and one day of test. In the acquisition, mice were performed three trials per day, for five consecutive days, to find a hidden black box located behind one of the 22 holes around the platform. A bright light source was placed above the platform to force the mice to explore the hidden box. The latency to the box was recorded for up to 180 s. Each mouse was allowed to remain in the box for 30 s and then moved from the maze to its home cage. If the mouse did not find the platform within 180 s, it was manually placed on the platform and returned to its home cage after 30 s. The inter-trial interval for each mouse was at least 20 min. On day 6, each mouse was tested for 60 s. Data were recorded using a video tracking software (TopScan, CleverSys, Inc.). The mean latency (in seconds) of the three trials was calculated for each day of trials. The latency to the hidden box and the number of sniffs to non-box holes (wrong sniffs) were calculated for the test trial.

**Y-maze test:** The Y-maze test was conducted to evaluate the short term spatial working memory of the mice, as previously described. The test contains two 5-min stages with an interval of 2 h between evaluation periods. During the first stage, the novel arm was blocked by a black baffle, allowing mice to only move in the other two arms. During the second stage, the novel arm was open, and mice could freely move throughout the three arms. The percentage of time traveled in each arm, number of entries into each arm, and travelling speed were calculated.

**Open field test:** Mice were carried to the behavior room to habituate at least 2 hours before starting the test. Mice were then placed into the open field arena (made of opaque blue plastic material, 50 cm × 50 cm) and allowed to explore the arena for 5 min. Total distance (cm) and percentage of time spent in the centre were quantified using video tracking software (TopScan, CleverSys, Inc.).

**Elevated plus maze test:** The elevated plus maze consisted of four arms (50 cm × 10 cm) and a central square (10 cm × 10 cm) connecting these arms, which was elevated 100 cm above the floor. Two opposite arms were open, while the remaining opposite arms were closed with 40 cm high walls. Each mouse was placed into the maze center facing an open arm, and left to freely explore the apparatus for 5 min. Arm entry was defined as entering an arm with all four paws. The open arm duration and entries were calculated.

**Tissue collection and processing**

Mice were anaesthed by i.p. injection of ketamine (80 mg/kg) and xylazine (8 mg/kg). Mice were then transcardially perfused with ice-cold PBS. For immunostaining, the brain and lungs were collected and drop-fixed in 4% PFA. Fixed brain were then washed with PBS, dehydrated with graded ethanol solutions and then embedded in paraffin or sucrose solution and then embedded in OCT compound respectively. Fixed and embedded brains or lungs were sliced (5 μm thick sections for paraffin embedded tissues or 15 μm for OCT embedded tissues) with a cryostat (Leica). For other experiments, brains, lung, meninges, spleens were dissected and flash frozen in liquid nitrogen and stored at −80 °C until analysis.

**Cell culture and treatment**

**Primary neurons**: The cortex of pregnant 16-19 d mice was removed and separated from meninges. Tissue were dissociated with 0.25% trypase at 37 °C and terminated by Neurobasal. Cells were resuspended with Neurobasal containing 1% B27, 0.5mM glutamine and 1% Penicillin-Streptomycin Solution and plated on 6-well plates. The culture medium was changed every 3 d and the cells were used at 6-7 d.

**Cell lines**: Murine neuronal lines N2a and human neuronal lines SH-SY5Y were purchased from ATCC. Cells were cultured in DMEM with 10% FBS and 1% Penicillin-Streptomycin at 37 ℃ with 5 % $CO_2$.

**Cell transfection:** siRNA targeting hSOCS1 or negative control (NC) siRNA was transfected in SH-SY5Y cell lines using Lipofectamine 2000 reagent in OPTI-MEM reduced serum medium according to the manufacturer's instructions.

**IL-35 treatment:** mouse IL-35 (Chimerigen Laboratories, CHI-MF-11135) or human IL-35 (Peprotech, 200-37) was added to culture medium at the concentration of 100 μg/ml. The cells were collected 48 h later expected for specific experiments.

**Immunohistochemistry**

Mouse brain or lung sections embedded OCT were subjected to a heat-induced antigen retrieval step with 10 mM citrate buffer for 15 min. For paraffin tissues, sections were subjected to the same antigen retrieval step after deparaffinization. For immunofluorescence staining, tissue was incubated in PBS and 0.3% Triton X-100 containing 5% bovine serum albumin (BSA) for 1 h at room temperature, and then incubated with appropriate dilutions of primary antibodies (Supplementary table 2) overnight at 4 °C. Appropriate donkey Alexa Fluor 488, 594 or 647 anti-rat, -goat, -rabbit or -mouse IgG antibodies (Thermo Fisher Scientific, 1:1000) were incubated for 1 or 2 h at room temperature in PBS. Then, DAPI was used to label nucleus following mounting with glass coverslips. Preparations were stored at 4 °C for no more than one week until images were acquired either using a confocal microscope (LSM 710 Laser Scanning Confocal Microscope, Zeiss). Quantitative analysis was performed on the acquired images using Fiji software.

**Flow cytometry**

Mice were injected i.p. with euthasol solution and were then transcardially perfused with ice-cold PBS. The meninges or lungs were dissected with fine forceps and digested in RPMI-1640 medium with 1.4 U/ml collagenase VIII (Sigma-Aldrich, C2139) and 1 mg/ml DNase I (Sigma-Aldrich, D4513) at 37 ℃ for 15 or 30 min, respectively. The brains and spleens were removed and smashed in PBS containing 2% FBS. The smashed brains were centrifuged with 30/70% percoll and single cell suspension was collected for following experiments. Erythrocyte lysis were conducted when needed. The cell pellets were washed, resuspended in ice-cold fluorescence-activated cell sorting (FACS) buffer (pH 7.4; 0.1 M PBS; 1 mM EDTA and 1% BSA). Cell viability was determined by using BD Horizon™ Fixable Viability Stain Reagents following the manufacturer's instructions. After an incubation period of 30 min at 4 °C, cells were washed and fixed in 1% PFA in PBS. Then, the cells were stained for extracellular markers with antibodies. Fluorescence data were collected with a FACS verse Cytometer (BD Bioscience) and analysed using FlowJo software (Tree Star, Inc.). In brief, singlets were gated using the height, area and the pulse width of the forward and side scatter and then viable cells were selected as FVS. Cells were then gated for the appropriate cell-type markers. An aliquot of unstained cells of each sample was counted using Automated Cell Counter (Merck) to provide accurate counts.

**Sorting of pulmonary B cells**

To obtain a suspension of pulmonary B cells using MACS, mice were euthanized by i.p. injection of euthasol and transcardially perfused with ice-cold PBS with heparin. The lungs were quickly collected and digested in RPMI-1640 medium with 1.4 U/ml collagenase VIII and 1 mg/ml DNase I for 30 min at 37 ℃. Individual samples with erythrocyte lysis were obtained after filtration through a 70-μm nylon-mesh cell strainer. Cell suspensions were then pelleted, resuspended in ice-cold MACS buffer containing anti-B220 microbeads (Miltenyi Biotec, 130-049-501) and incubated for 30 min at 4 °C. Cells were rewashed and resuspended in ice-cold MACS buffer, then passed through MACS column and collected for the following experiments.

**B cells tracking experiments**

B cells sorted from the lung of 3-month-old WT or 5xFAD mice were firstly labeled by PKH26 Red Fluorescent Cell Linker Kits (Sigma-Aldrich, MINI26) according to the instructions. Then, 3-month-old 5xFAD mice were infused with $2\times10^5$ cells per mouse via retro-orbital injection. Twenty-four hours after transplantation, IVIS Spectrum In Vivo Imaging System (Perkin-Elmer) was used to detected the distribution of labeled cells. Finally, the mice were sacrificed for the following immunofluorescence analysis.

**Western blotting**

For Western blot analyses, the homogenized protein samples of brain or lung were loaded onto 10–15% Tris/tricine SDS gels, and transferred to PVDF membranes. After blocking for 1 h in 5% nonfat milk/TBST, the membranes were incubated at 4℃ overnight with primary antibodies (Supplementary table 2). Horseradish peroxidase-conjugated secondary antibodies (Vector Laboratories, USA) were used, and bands were visualized using ECL plus detection system. GAPDH was used as an internal reference for protein loading and transfer efficiency. Four mice per group in duplicate experiments were averaged to provide a mean value for each group.

**RNA extraction and sequencing**

For total RNA extraction, the tissues or cells was immersed in the appropriate volume of Trizol, immediately snap-frozen in liquid nitrogen and stored at −80 °C until further use. After defrosting on ice, samples were mechanically dissociated in extraction buffer and RNA was isolated using the kit components according to the manufacturer’s instructions (RNAiso Plus, Takara, 9109). The Takara RNA Library Prep Kit was used for cDNA library preparation from total RNA samples. Relative expression of mRNA for the target genes was performed by the comparative CT (△△CT) method using GAPDH as control reference genes. The primers were listed in Supplementary table 3.

The RNA sequence of pulmonary B cells was performed by Cloud-seq. The raw sequencing reads (FASTQ files) were first chastity filtered, which removed any clusters that have a higher than expected intensity of the called base compared to other bases. The quality of the reads was then evaluated using FastQC, and after passing quality control, the expression of the transcripts was quantified against the UCSC mm 10 genome using Salmon. These transcript abundances were imported into R and summarized with tximport, and then edgeR was used to normalize the raw counts, and perform exploratory analysis and differential expression analysis. The *P* values from the differential expression analysis were corrected for multiple hypothesis testing with the Benjamini–Hochberg false-discovery rate procedure (adjusted *P* value). Functional enrichment of differential expressed genes, using gene sets from Gene Ontology (GO), Kyoto Encyclopedia of Genes and Genomes (KEGG) or gene-set enrichment analysis (GSEA), was determined with Fisher’s exact test as implemented in the cluster Profiler Bioconductor package. Heat maps of the differential expressed genes and enriched gene sets were generated with the R package “pheatmap”. Normalized counts of selected transcripts were used to calculate the fold change relative to respective controls.

**ELISA Analysis**

Frontal cortex samples were homogenized and sonicated in ice-cold TBS buffer containing 0.5 mM PMSF, 0.5 mM benzamidine, 1.0 mM DTT and 1.0 mM EDTA, followed by centrifugation at 12000 × g for 30 min. Supernatants were set aside for measurements of IL-35. Pellets were re-suspended and further homogenized in 70 % formic acid (equal volume of TBS), then centrifuged at 12000 × g for 1 h. The above indexes were quantified with ELISA kits from Biolegend according the manufacturer’s instructions.

**Luciferase Assay**

The promoter of hBACE1 gene was constructed into pGL3 plasmids and transfected into SH-SY5Y cells with Lipofectamine 2000 reagent (Invitrogen, #52887). Human IL-35 (Peprotech, 200-37) was added after 6-8 h, and the cells were collected 24 h later. According to the instructions of the promega luciferase detection kit (Promega, E1910), the cells were washed twice with PBS, diluted with lysis buffer and shaked in a horizontal mixer for 15 min. The fluorescence was detected using the GloMax® 20/20 Luminometer.

**DNA pulldown and mass spectrometry**

Nuclear extracts were prepared as described above for EMSAs. 10 μg of biotinylated double-stranded DNA oligonucleotides was immobilized on 0.5 mg of Dynabeads M-280 Streptavidin (Thermo Fisher, 11205D) and incubated overnight with 100 μl of nuclear extract and 1× binding buffer (10 mM HEPES pH 7.9, 4% glycerol, 60 mM KCl, and 1 mM MgCl2). Beads were washed tiwce with wash buffer (60 mM KCl, 1 mM MgCl2, and 0.2% NP-40) and twice with wash buffer without NP-40. After streptavidin bead pulldown, proteins were digested on bead and prepared for mass spectrometry. Data were quantified and searched against the UniProt mouse database (July 2016) using MaxQuant (version 1.5.0.30).

**BACE1 enzyme activity analysis**

BACE1 enzyme activity detection was performed according the manufacturer's instructions (β-Secretase Activity Fluorometric Assay Kit, Sigma-Aldrich, MAK237). Briefly, the lysis of the cell was centrifuged at 10,000 g for 5 min, and a portion was used for protein quantitative experiment with BCA method. 50 μL of cell lysis was used, and reaction buffer and substrate were added subsequently. The fluorescence of each sample was detected using a full-wavelength microplate reader.

**Statistical analysis and reproducibility**

The data analysts were blind to the identity of the experimental group. Statistical tests for each figure were justified to be appropriate using Prism 6.0 (GraphPad Software, Inc.). One-way ANOVA, with Bonferroni's post hoc test or Holm-Sidak's post hoc test, was used to compare three independent groups. Two-group comparisons were made using two-tailed unpaired student's t-tests. For comparisons of multiple factors, two-way ANOVA with Bonferroni's post hoc test was used. Repeated-measures ANOVA with Bonferroni's post hoc test was used for day versus treatment comparisons with repeated observations. Data are always presented as mean ± s.e.m.

### Supplementary Figures and Figure legends:

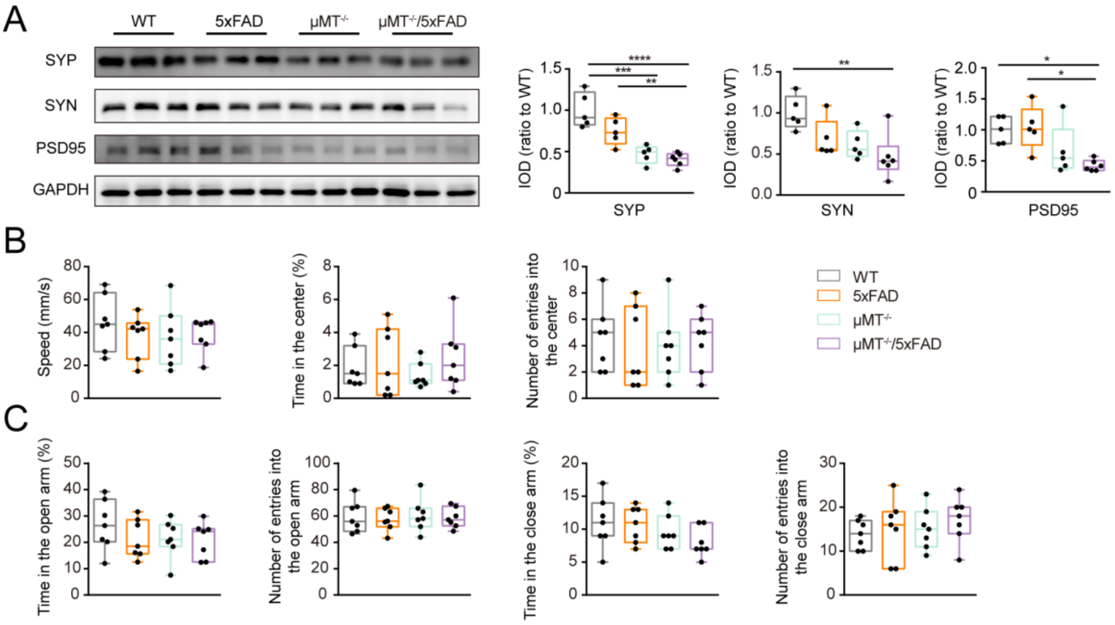


**Fig. S1. (Related to Fig. 1) B lymphocytes deletion does not result in the anxiety-like behavior of 5xFAD mice at 3 months old.** (**A**) Representative bands and quantification of SYP, SYN and PSD95 in the frontal cortex. n = 5-6 per group. (**B**) Quantification of the speed, the percentage of time in the central area and the number of times in the central area of 3-month-old WT, 5xFAD, $\mu MT^{-/-}$ and $\mu MT^{-/-}$/5xFAD mice in the open field experiment. (**C**) Quantification of the percentage of time in the open arm, the number of times in the open arm and the percentage of time in the closed arm. n = 7 per group. *$P < 0.05$, **$P < 0.01$, ***$P < 0.001$, ****$P < 0.0001$. Data in **A** to **C** ANOVA with Bonferroni's post hoc test.

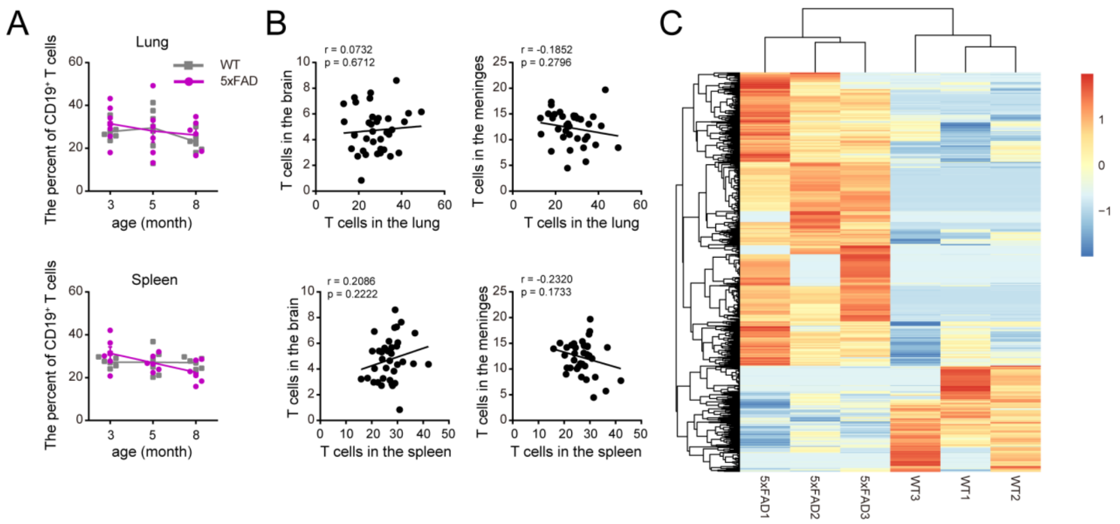


**Fig. S2. (Related to Fig. 2) Changes of lymphocytes between the lungs and spleen of 3-, 5-and 8-month-old 5xFAD mice.** (**A**) Percentage of T lymphocytes in the lungs (upper panel) and spleen (lower panel) of 3-, 5- and 8-month-old 5xFAD mice. n = 6 per group. (**B**) Correlation analysis of T lymphocytes between the lungs and brain, and meninges, respectively (upper panel), between the spleens and brain, and meninges (lower panel). (**C**) Heat map of different up- and down-regulated genes in the pulmonary B cells of 3-month-old WT and 5xFAD mice. The color scale is the r-log-transformed values across samples. n = 3 per group Data in **A** by ANOVA with Bonferroni’s post hoc test, **B** by Pearson correlation coefficient and **C** by Fisher's exact test.

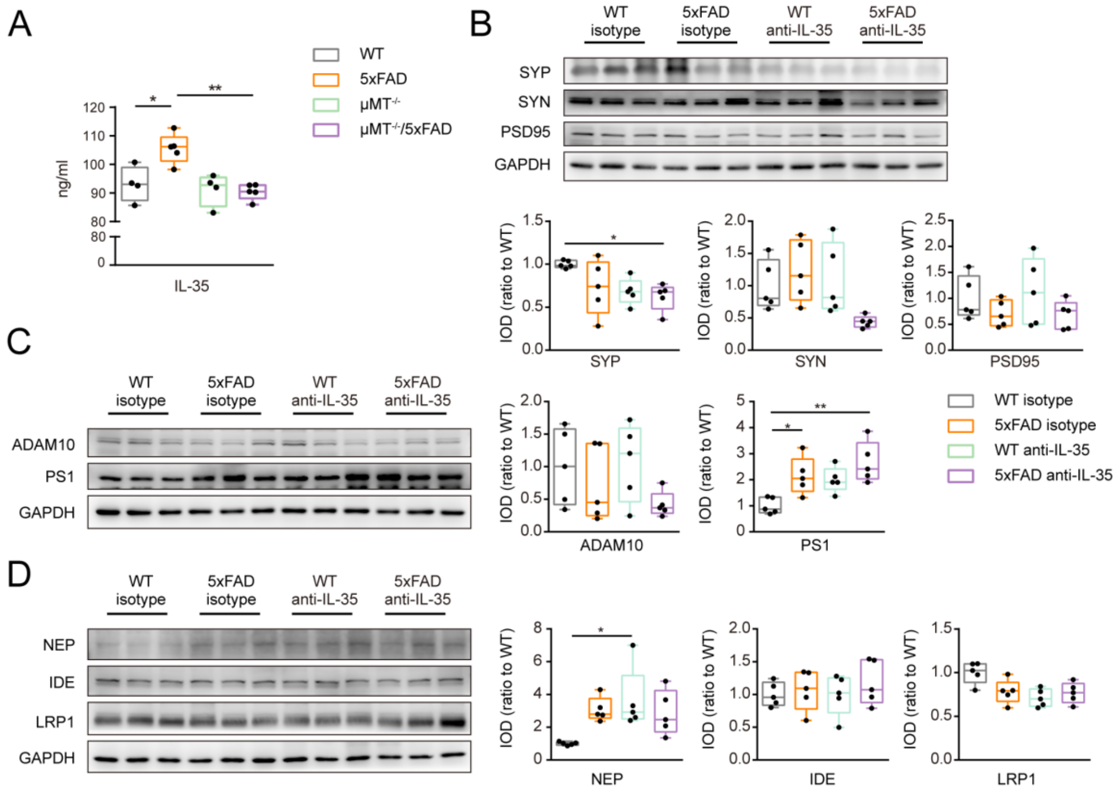


**Fig. S3. (Related to Fig. 3 and Fig. 4) Synapses and Aβ metabolism related proteins in the frontal cortex of 3-month-old 5xFAD mice without B cells or after neutralization of IL-35.** (**A**) ELISA for IL-35 in the frontal cortex of WT, 5xFAD, *μMT*$^{-/-}$ and *μMT*$^{-/-}$/5xFAD mice. n = 4-5 per group. (**B**) Representative bands and quantification of western blot for SYP, SYN and PSD95 in the frontal cortex of the above genotype mice. n = 5 per group. (**C** and **D**) Representative bands and quantification of western blot for ADAM10, PS1, NEP, IDE, LRP1 in the PFC of 5xFAD mice after neutralization of IL-35. n = 5 per group. **P* < 0.05, ***P* < 0.01, ****P* < 0.001, *****P* < 0.0001. Data in **A** to **D are analyzed by** ANOVA with Bonferroni's post hoc test.

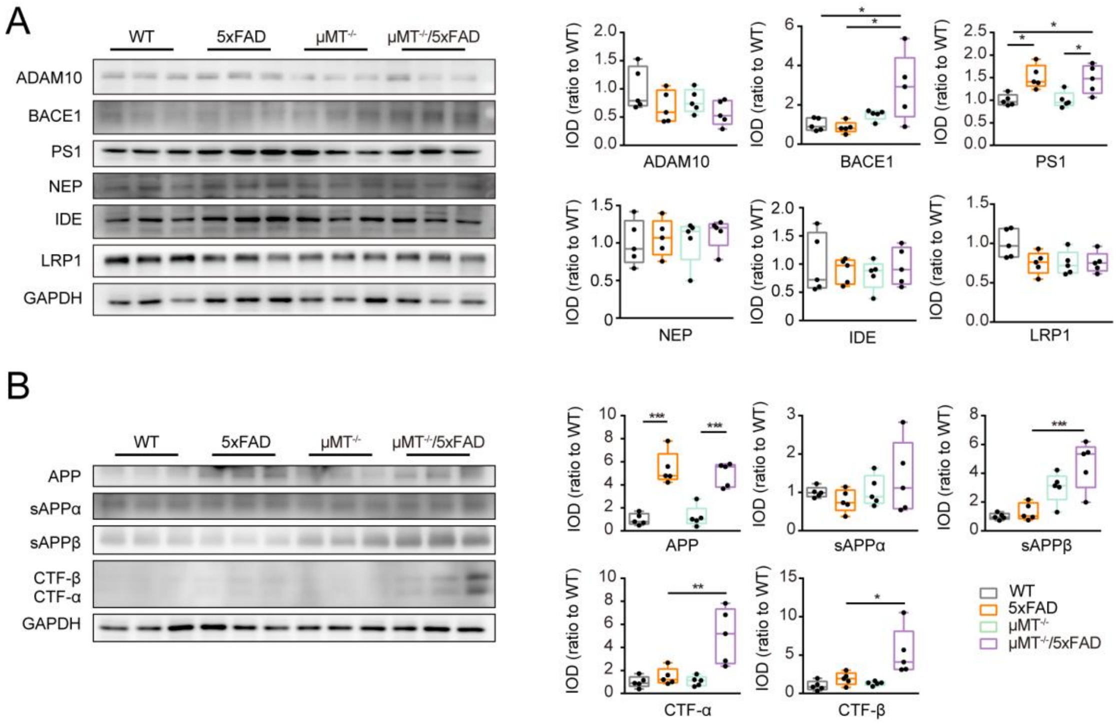


**Fig. S4. (Related to Fig. 4) The expression levels of Aβ production and degradation related proteins in the frontal cortex of 3-month-old 5xFAD mice and *μMT*$^{-/-}$/5xFAD mice.** (**A**) Representative bands and quantification of Western blot for ADAM10, BACE1, PS1, NEP, IDE and LRP1 in the PFC of WT, 5xFAD, *μMT*$^{-/-}$ and *μMT*$^{-/-}$/5xFAD mice. n = 5 per group. (**B**) Representative bands and quantification of Western blot for APP, sAPPα, sAPPβ, CTF-α and CTF-β in the frontal cortex of the above genotype mice. n = 5 per group. *$P < 0.05$, **$P < 0.01$, ***$P < 0.001$, ****$P < 0.0001$. Data in **A**, **B are analyzed** ANOVA with Bonferroni's post hoc test.

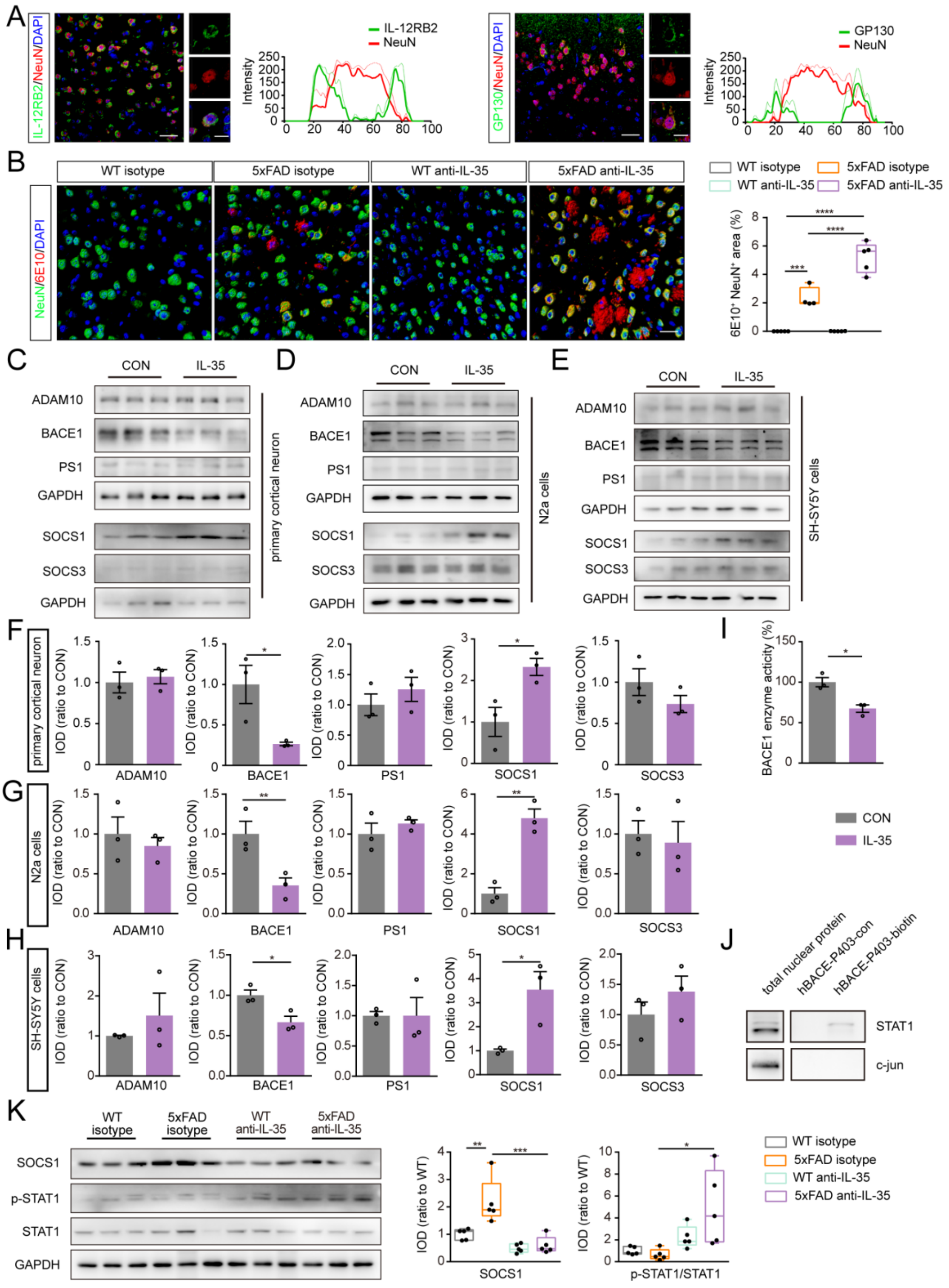

**Fig. S5. (Related to Fig. 4) IL-35 inhibits BACE1 expression by SOCS1 pathway.** (**A**) Representative images of IL-12RB2 or GP130 with NeuN respectively and analysis of their co-localization in the frontal cortex of 3-month-old mice, Scale bar = 30 μm. (**B**) Representative images and quantification of NeuN and 6E10 in the frontal cortex of 3-month-old mice after IL-35 neutralization. n = 4-5 per group. Scale bar = 30 μm. (**C** and **F**) Representative bands and quantification of Western blot for ADAM10, BACE1, PS1, SOCS1 and SOCS3 in the mouse primary neuron after IL-35 treatment. (**D** and **G**) Representative bands and quantification of western blot for ADAM10, BACE1, PS1, SOCS1 and SOCS3 in the N2a cell line after IL-35 treatment. (**E** and **H**) Representative bands and quantification of Western blot for ADAM10, BACE1, PS1, SOCS1 and SOCS3 in the SH-SY5Y cell line after IL-35 treatment. n = 3 per group. (**I**) Activity of BACE1 enzyme in primary cortical neurons after IL-35 treatment. n = 3 per group. (**J**) Representative bands of Western blot for STAT1 and c-jun after hBACE1-P403 pulldown in SH-SY5Y cells. (**K**) Representative bands and quantification of Western blot for SOCS1, p-STAT1 and STAT1 in 3-month-old mice after IL-35 neutralization. n = 5 per group. $*P < 0.05$, $**P < 0.01$, $***P < 0.001$, $****P < 0.0001$. Data in **B, K** by ANVOA with Bonferroni's post hoc test, others by Student's t-test.

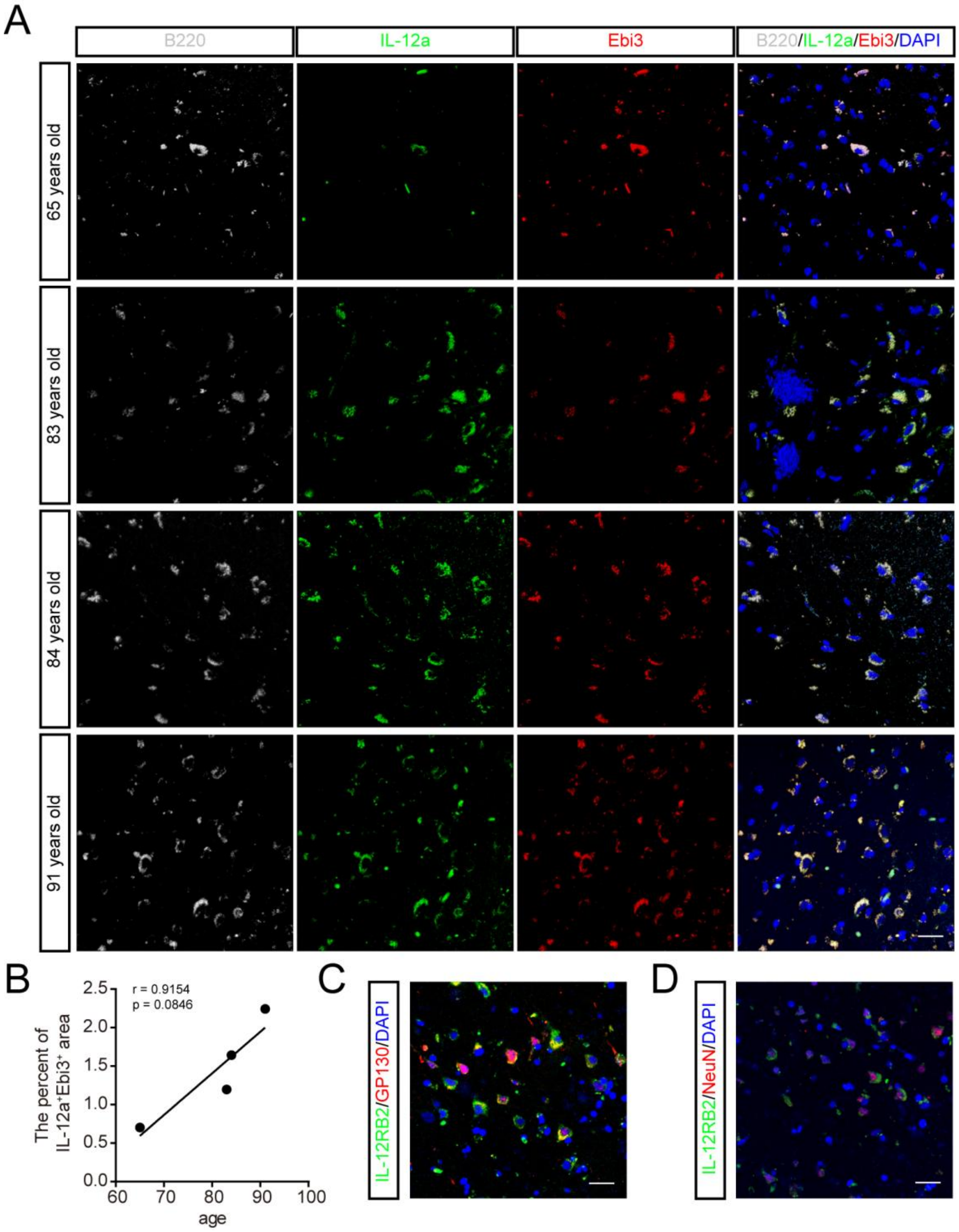


**Fig. S6 IL-35 expression of B lymphocytes and its receptors in postmortem brain.** (**A**) Representation images for IL-12a, Ebi3 and B220 in the human frontal cortex. (**B**) Correlation analysis of the age and the percent of IL-12a$^{+}$Ebi3$^{+}$ area in the cortex. n = 4. (**C**) Co-localization of IL-12RB2 and GP130in the human frontal neurons. (**D**) IL-12RB2 expression in the neurons of postmortem brain. Scale bar = 30 μm. Data in **B** was analyzed by Pearson Correlation Coefficient.

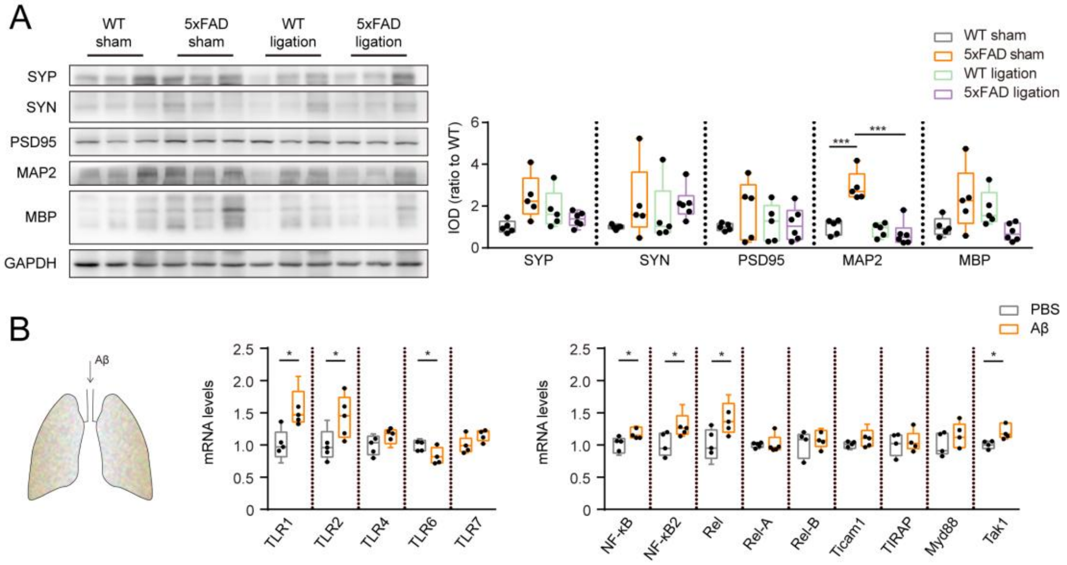


**Fig. S7. (Related to Fig. 5) Aβ induced the up-regulation of IL-35 in pulmonary B cells.** (**A**) Representative bands and quantification of western blot for SYP, SYN, PSD95, MAP2 and MBP in the lungs of WT and 5xFAD mice after dcLN ligation. n = 5-6 per group. (**B**) Q-PCR for toll-like receptors and NF-κB pathways relative genes in the pulmonary B cells after Aβ stimulation in 3-month-old WT mice. n = 5 per group. *$P < 0.05$, **$P < 0.01$, ***$P < 0.001$, ****$P < 0.0001$. Data in **A** ANOVA with Bonferroni's post hoc test, **B** Student's t-test.

**Supplementary table 1: The proteins showing significant interaction with the hBACE-P403**

| Gene name | Unique peptides | iBAQ | Gene name | Unique peptides | iBAQ |
|---|---|---|---|---|---|
| HNRNPU | 22 | 1440900 | DHX30 | 3 | 4401 |
| SF3B1 | 22 | 173920 | EIF3B | 3 | 10509 |
| MYH9 | 21 | 64441 | FAM120A | 3 | 14042 |
| TOP2A | 14 | 231020 | FUS | 3 | 48259 |
| SFPQ | 18 | 5260700 | GLG1 | 3 | 11998 |
| DHX9 | 16 | 163150 | HNRNPC | 3 | 86476 |
| TOP2B | 9 | 31314 | KIAA1429 | 3 | 17263 |
| HNRNPM | 14 | 132660 | LIMA1 | 3 | 21406 |
| RBM14 | 14 | 382730 | LMNA | 2 | 21422 |
| RBM25 | 14 | 216930 | LMNB1 | 2 | 24441 |
| SMARCA5 | 14 | 79395 | MYO1C | 3 | 10279 |
| MATR3 | 13 | 240720 | NOLC1 | 3 | 93701 |
| NONO | 11 | 839940 | NOP58 | 3 | 12966 |
| RRBP1 | 13 | 198650 | NUMA1 | 3 | 44439 |
| DDX21 | 12 | 398670 | PPP1R9B | 3 | 19248 |
| HNRNPA2B1 | 12 | 871940 | PRPF6 | 3 | 1781.6 |
| DDX17 | 7 | 162950 | PRPF8 | 3 | 3498.1 |
| EFTUD2 | 11 | 101940 | RBM15 | 3 | 11797 |
| KTN1 | 11 | 64793 | RBM27 | 2 | 13316 |
| PNN | 11 | 179350 | RPL11 | 3 | 52910 |
| PRPF40A | 11 | 301380 | RPL12 | 3 | 93175 |
| SF3A1 | 11 | 196260 | RPL13a | 3 | 722670 |
| SUPT16H | 11 | 100020 | RPL17 | 3 | 144900 |
| DDX5 | 6 | 57185 | RPL22L1 | 3 | 98082 |
| HNRNPK | 10 | 284980 | RPL34 | 3 | 458130 |
| YBX1 | 7 | 1526400 | RPL5 | 3 | 79607 |
| RPL7 | 9 | 928760 | RPS11 | 3 | 206930 |
| CAPRIN1 | 8 | 553140 | RPS17 | 3 | 112030 |
| DDX23 | 8 | 25919 | RPS25 | 3 | 546080 |
| DDX46 | 8 | 38225 | RPS5 | 3 | 108230 |
| HDLBP | 8 | 24703 | RPSA | 3 | 51340 |
| HNRNPUL2 | 8 | 62974 | SART1 | 3 | 10578 |
| RPL4 | 8 | 153480 | SEC63 | 3 | 20060 |
| SF3B3 | 8 | 44688 | SLTM | 3 | 29685 |
| STT3A | 8 | 193350 | SNRNP200 | 3 | 3760.4 |
| THRAP3 | 7 | 44340 | SNRNP70 | 3 | 39146 |
| UBTF | 8 | 35968 | SNRPN | 3 | 71955 |
| ATP5B | 7 | 214160 | SRSF11 | 3 | 98209 |
| DDX3X | 6 | 51039 | SRSF7 | 3 | 302470 |
| HNRNPA1 | 7 | 325220 | TUBB | 3 | 43709 |

| | | | | | |
|---|---|---|---|---|---|
| NCL | 7 | 547660 | UBAP2L | 3 | 13128 |
| RBMX | 7 | 133980 | ZC3H18 | 3 | 25420 |
| RPS4X | 7 | 107760 | ABCF2 | 2 | 6152.9 |
| U2SURP | 7 | 58703 | ATAD3B | 2 | 10009 |
| G3BP1 | 5 | 65255 | ATP2A2 | 2 | 3366.5 |
| NOP2 | 6 | 177830 | BAZ1A | 2 | 2643.6 |
| RPL32 | 6 | 341510 | CANX | 2 | 17917 |
| RRP12 | 6 | 16204 | CCDC47 | 2 | 12657 |
| SF3B2 | 6 | 95211 | CCNK | 2 | 21551 |
| SND1 | 6 | 16807 | CHTOP | 2 | 93282 |
| YTHDC1 | 6 | 66235 | COPA | 2 | 243.05 |
| SLC25A5 | 3 | 315500 | CPSF1 | 2 | 2176 |
| ASPH | 5 | 63703 | DHX37 | 2 | 6977.8 |
| G3BP2 | 4 | 85634 | EIF3A | 2 | 0 |
| RPL23A | 5 | 349300 | EIF3C | 2 | 209.09 |
| RPS3A | 5 | 352830 | EWSR1 | 2 | 80264 |
| RPS6 | 5 | 401620 | FBL | 2 | 27732 |
| SEC61A1 | 5 | 107720 | FTSJ3 | 2 | 14603 |
| SPECC1L-ADORA2A | 5 | 14535 | GTPBP4 | 2 | 4765.3 |
| SRSF1 | 5 | 229330 | HNRNPAB | 2 | 41561 |
| SRSF3 | 5 | 394380 | HNRNPL | 2 | 18926 |
| ZFR | 5 | 35564 | HSP90AB1 | 2 | 7776.1 |
| CDK11B | 4 | 14132 | IGF2BP3 | 2 | 11237 |
| CWC22 | 4 | 19849 | ILF3 | 2 | 8526 |
| DDX1 | 4 | 16816 | IMMT | 2 | 6016.2 |
| HP1BP3 | 4 | 33285 | INTS3 | 2 | 4431.8 |
| PHB | 4 | 470890 | IPO7 | 2 | 2085.7 |
| PPP1R12A | 4 | 20478 | LUC7L3 | 2 | 51848 |
| PRPF38B | 4 | 209740 | MPRIP | 2 | 4425.9 |
| HNRNPA3 | 2 | 269270 | MYBBP1A | 2 | 115660 |
| PSPC1 | 4 | 121950 | NEXN | 2 | 53617 |
| RBM6 | 4 | 11619 | NOMO3 | 2 | 10774 |
| RPS16 | 4 | 53319 | NOP14 | 2 | 2941.4 |
| RPS2 | 4 | 153040 | PARP1 | 2 | 1278.9 |
| SMC2 | 4 | 11234 | PAXBP1 | 2 | 11893 |
| SRRM1 | 4 | 63756 | PDS5B | 2 | 1730.7 |
| SRRT | 4 | 32763 | PUF60 | 2 | 19677 |
| SRSF2 | 4 | 2129300 | RALY | 2 | 54821 |
| SSRP1 | 4 | 31480 | RBM28 | 2 | 15855 |
| YBX3 | 2 | 197990 | RBM3 | 2 | 139470 |
| RPL7A | 10 | 1707700 | RPL10A | 2 | 26440 |
| BCLAF1 | 3 | 25726 | RPL18A | 2 | 269240 |
| RPS3 | 11 | 355780 | RPL27 | 2 | 2419000 |

| | | | | | |
|---|---|---|---|---|---|
| ACTG1 | 6 | 823690 | RPL27A | 2 | 170230 |
| RPL6 | 6 | 1015500 | RPL31 | 2 | 201180 |
| RPL18 | 5 | 1799000 | RPL36A | 2 | 340300 |
| RPS8 | 6 | 985120 | RPL9 | 2 | 39075 |
| HSPA8 | 3 | 34141 | RPS19 | 2 | 32975 |
| ATP5A1 | 6 | 47159 | RPS20 | 2 | 80242 |
| VIM | 17 | 629400 | RPS27L | 2 | 116180 |
| RPL15 | 8 | 1183400 | RPS7 | 2 | 62870 |
| STAT1 | 10 | 69164 | SAFB | 2 | 20673 |
| RPL3 | 8 | 373790 | SCAF4 | 2 | 3061.4 |
| RPL13 | 5 | 1483700 | SERBP1 | 2 | 24924 |
| HIST1H1E | 10 | 7643800 | SF3A2 | 2 | 55297 |
| H1F0 | 5 | 569210 | SLC25A3 | 2 | 228090 |
| SRSF6 | 3 | 618390 | STT3B | 2 | 3593.2 |
| ACIN1 | 3 | 18395 | TARDBP | 2 | 51865 |
| AEBP1 | 3 | 13788 | TRIM28 | 2 | 22597 |
| ARHGEF2 | 3 | 5777.4 | U2AF1L5 | 2 | 98860 |
| CDC5L | 3 | 11795 | UPF1 | 2 | 7050 |
| CHERP | 3 | 25735 | WDR33 | 2 | 23457 |

**Supplementary table 2: antibody information**

| | | | | Dilution | | |
|---|---|---|---|---|---|---|
| **Antibodies** | **Source** | **Catalog number** | **Host species&clone** | **WB** | **IHC** | **Flow** |
| CD45-FITC | eBioscience | 11-0451 | rat monoclonal | | | 1:200 |
| CD3-eF450 | eBioscience | 48-0032 | rat monoclonal | | | 1:200 |
| CD19-PE-CY7 | eBioscience | 25-0193 | rat monoclonal | | | 1:200 |
| 6E10 | biolegend | 803001 | mouse monoclonal | 1:1000 | 1:500 | |
| GFAP | millipore | MAB360 | mouse monoclonal | | 1:500 | |
| GFAP | abcam | ab4674 | chicken polyclonal | | 1:500 | |
| Iba-1 | wako | 019-19741 | rabbit polyclonal | | 1:500 | |
| Iba-1 | abcam | ab5076 | goat polyclonal | | 1:500 | |
| SYP | millipore | MAB5258-I | mouse monoclonal | 1:1000 | | |
| SYN | abcam | ab64581 | rabbit polyclonal | 1:1000 | | |
| PSD95 | abcam | ab18258 | rabbit polyclonal | 1:1000 | | |
| GAPDH | proteintech | 60004-1-Ig | mouse monoclonal | 1:3000 | | |
| IL-12a | abcam | ab131039 | rabbit monoclonal | 1:1000 | | |
| IL-12a | R&D | MAB1570 | mouse monoclonal | | 1:250 | |
| Ebi3 | abcam | ab124694 | rabbit monoclonal | 1:1000 | | |
| Ebi3 | santa cruz | sc-166158 | mouse monoclonal | | 1:200 | |
| B220 | eBioscience | 14-0452 | rat monoclonal | | 1:50 | |
| ADAM10 | millipore | AB19026 | rabbit polyclonal | 1:1000 | | |
| BACE1 | millipore | MAB5308 | mouse monoclonal | 1:1000 | 1:200 | |
| BACE1 | CST | 5606 | rabbit monoclonal | 1:1000 | | |
| PS1 | sigma | PRS4203 | rabbit polyclonal | 1:1000 | | |
| sAPPα | IBL | 11088 | mouse monoclonal | 1:1000 | | |
| sAPPβ | biolegend | 813401 | rabbit polyclonal | 1:1000 | | |
| APP | sigma | sab4300464 | rabbit polyclonal | 1:1000 | | |
| NEP | millipore | AB5458 | rabbit polyclonal | 1:1000 | | |
| IDE | abcam | ab32216 | rabbit polyclonal | 1:1000 | | |
| LRP1 | abcam | ab92544 | rabbit monoclonal | 1:1000 | | |
| IL-12RB2 | LSBio | LS-C29482 9 | rabbit polyclonal | 1:1000 | 1:250 | |
| GP130 | Santa cruz | sc-376280 | mouse monoclonal | 1:1000 | 1:200 | |
| NenN | millipore | MAB377 | mouse monoclonal | | 1:200 | |
| NenN | abcam | Ab177487 | rabbit monoclonal | | 1:800 | |
| SOCS1 | CST | 3950 | rabbit polyclonal | 1:1000 | | |
| SOCS3 | proteintech | 14025-1-AP | rabbit polyclonal | 1:1000 | | |
| STAT1 | CST | 14994 | rabbit monoclonal | 1:1000 | 1:500 | |
| P-STAT1(Tyr701 ) | CST | 9167 | rabbit monoclonal | 1:1000 | | |
| BAFF | R&D | MAB1357 | rat monoclonal | 1:1000 | | |
| MAP2 | millipore | AB5622 | rabbit polyclonal | 1:1000 | | |
| MBP | abcam | ab7349 | rat monoclonal | 1:1000 | | |

**Supplementary table 3: qPCR primer information**

| Primer | Primer sequence (5' to 3' ) |
|---|---|
| IL-12a-F | CATCGATGAGCTGATGCAGT |
| IL-12a-R | CAGATAGCCCATCACCCTGT |
| Ebi3-F | TGCTCTTCCTGTCACTTGCC |
| Ebi3-R | CGGGATACCGAGAAGCATGG |
| IL-10-F | GCCCTTTGCTATGGTGTCCTTTC |
| IL-10-R | TCCCTGGTTTCTCTTCCCAAGAC |
| TGF-β-F | TTGCTTCAGCTCCACAGAGA |
| TGF-β-R | TGGTTGTAGAGGGCAAGGAC |
| IL-6-F | GGAGCCCACCAAGAACGATA |
| IL-6-R | AGACAGGTCTGTTGGGAGTG |
| TNF-α-F | CAGGCGGTGCCTATGTCTC |
| TNF-α-R | CGATCACCCCGAAGTTCAGTAG |
| BAFF-F | ACACTGCCCAACAATTCCTG |
| BAFF-R | TCGTCTCCGTTGCGTGAAATC |
| TLR1-F | TGAGGGTCCTGATAATGTCCTAC |
| TLR1-R | AGAGGTCCAAATGCTTGAGGC |
| TLR2-F | GCAAACGCTGTTCTGCTCAG |
| TLR2-R | AGGCGTCTCCCTCTATTGTATT |
| TLR4-F | GCTTTCACCTCTGCCTTCAC |
| TLR4-R | GAAACTGCCATGTTTGAGCA |
| TLR6-F | TGAGCCAAGACAGAAAACCCA |
| TLR6-R | GGGACATGAGTAAGGTTCCTGTT |
| TLR7-F | ATGTGGACACGGAAGAGACAA |
| TLR7-R | GGTAAGGGTAAGATTGGTGGTG |
| NF-κB-F | TGGCTTTGCAAACCTGGGAA |
| NF-κB-R | AATACACGCCTCTGTCATCCGT |
| NF-κB2-F | GGCCGGAAGACCTATCCTACT |
| NF-κB2-R | CTACAGACACAGCGCACACT |
| Rel-F | AGAGGGGAATGCGGTTTAGAT |
| Rel-R | TTCTGGTCCAAATTCTGCTTCAT |
| Rel-A-F | GCCCAGACCGCAGTATCC |
| Rel-A-R | GTCCCGCACTGTCACCTG |
| Rel-B-F | CCGTACCTGGTCATCACAGAG |
| Rel-B-R | CAGTCTCGAAGCTCGATGGC |
| Ticam1-F | ACCTTCTGCGAGGATTTCCA |
| Ticam1-R | CGACAGTCGAAGTTGGAGGT |
| TIRAP-F | CCTCCTCCACTCCGTCCAA |
| TIRAP-R | CTTTCCTGGGAGATCGGCAT |
| Myd88-F | TCATGTTCTCCATACCCTTGGT |
| Myd88-R | AAACTGCGAGTGGGGTCAG |
| Tak1-F | GTCATCCAGCCCTAGTGTCAGAAT |
| Tak1-R | TTCTTTGGAGTTTGGGCACG |

| | |
|---|---|
| GAPDH-F | AGGTCGGTGTGAACGGATTTG |
| GAPDH-R | GGGGTCGTTGATGGCAACA |
